# abcFISH enables multiplexed, single-molecule visualization of circular RNA spatial heterogeneity

**DOI:** 10.1101/2025.11.16.688740

**Authors:** Yu-Xin Liu, Bo-Wen Jiang, Kai-Ming Liu, Li Yang, Ling-Ling Chen

## Abstract

Functional RNAs often exhibit distinct subcellular localization, but studying circular RNA (circRNA) localization has been challenging due to their extensive sequence overlap with linear mRNAs. We developed <u>a</u>mplicon-<u>b</u>ased <u>c</u>ircular RNA <u>f</u>luorescence *in <u>s</u>itu* <u>h</u>ybridization (abcFISH), a method that employs an optimized rolling circle amplification (RCA) strategy targeting back-splicing junction (BSJ) sites for robust, high-specificity, single-molecule circRNA imaging. abcFISH enables quantitative, multiplexed imaging in cells and tissues, allowing us to reveal alternative circularization patterns from the single locus; uncover cell-type-specific *circPOLR2A(9,10)* expression related to a combinatorial effect of RNA-binding proteins; map the distinct spatial distribution patterns of multiple circRNAs in neurons and brain tissues; and show the functional interplay of circRNA *Cdr1as* and lncRNA *Cyrano* co-localization. Furthermore, we applied abcFISH to validate the efficacy of therapeutic double-stranded circRNA aptamers (ds-cRNAs), simultaneously visualizing AAV-delivered ds-cRNAs and a consequent reduction in astrocyte infiltration around transduced cells. Collectively, abcFISH establishes a robust and user-friendly toolkit for deciphering circRNA localization and function *in vivo*.

## Introduction

Circular RNAs (circRNAs) share the same primary sequence as their linear counterparts, with exception of the back-splicing junction (BSJ), where a downstream 5’ splice donor is joined to an upstream 3’ splice acceptor^1^. Consequently, circRNA annotation, validation, and functional studies rely on developing appropriate methods to discriminate sequences spanning the unique BSJ^2^. As with other regulatory RNAs, the functions of circRNAs are closely linked to their abundance and subcellular localization within cells^3^. CircRNAs are dynamically expressed across different developmental and differentiation stages^4–6^. While computational analysis of the BSJ-spanning reads from high-throughput RNA sequencing (RNA-seq) datasets enables circRNA quantification, discrepancies exist between algorithms, underscoring the need for careful circRNA annotation^7, 8^. Further quantifying the cellular circRNA copy number can be achieved by measuring band intensity on Northern blotting gels^9^, or by comparing Ct values to their relative standard curves^10^, respectively. While subcellular localization can be assessed through bulk cell fractionation followed by qPCR^6^, this approach is often compromised by imperfect separation and lack of single-cell information^1^.

RNA *in situ* hybridization (ISH) provides a direct method for determining the circRNA subcellular localization but faces several challenges. Early studies used long (∼100 nt) digoxigenin (DIG)-labeled antisense probes spanning BSJ to detect abundant circRNAs in mouse brain sections^11^. However, these probes could also hybridize with cognate linear mRNAs. Single-molecule FISH (smFISH), employing a set of ∼32-48 labeled DNA oligos, has been used to detect *CDR1as*, a human circRNA, antisense to the cerebellar degeneration-related protein 1 transcript with the length of approximately 1,400 nt^12, 13, 14^. Yet, such probes may similarly detect its linear parental transcript *LINC00632*^15^. Although a two-fluorophore smFISH strategy, referred as circFISH, could distinguish circular from its cognate linear RNAs^16^, this method is limited to large circRNAs and remains ineffective for the majority that are relatively small with low abundance^17, 18^.

Imaging circRNAs via their BSJ sites overcomes the length limitation inherent to smFISH. Signal amplification-based methods, such as the branched DNA system of ViewRNA^19, 20^ and the “Z” probe technology of BaseScope^21^ are available but involve proprietary, multi-step procedures that make them costly. Alternatively, rolling-circle amplification (RCA) FISH offers robust signal enhancement^22^ and has been applied to *in situ* sequencing^23, 24^, ribosome-bound mRNA imaging^25^ and glycoRNA detection^26^. Inspired by this versatility, we sought to explore its potential for imaging circRNAs. Notably, a previous study applied RCA for circRNA imaging using SplintR ligase^27^. However, this approach yielded limited sensitivity, likely due to the ligation efficiency constraints at the BSJ nick. Here, we applied T4 DNA ligase with a different RCA probe design, achieving substantially higher sensitivity and lower cost for circRNA detection.

Inspired by this versatility, we designed RCA probes targeting circRNA BSJ sites, establishing a method we termed amplicon-based circular RNA fluorescence in situ hybridization (abcFISH). abcFISH exhibited high specificity, high sensitivity and low cost over existing techniques^16, 27–29^, enabling multiplexed and quantitative imaging of circRNAs in single cells and detection of alternative circRNA isoforms sharing the same upstream splice acceptor. Applied to tissue sections, it revealed heterogeneous expression of human *circPOLR2A(9,10)* in a knock-in (KI) mouse model^30^ and distinct subregional and subcellular localization of endogenous circRNAs, such as *Cdr1as* and *circAnks1b(5,6,7,8)* in the hippocampus with neuronal enrichment. abcFISH also evaluated the functional interplay between *Cdr1as* and the long noncoding RNA *Cyrano*^31^ within the soma, detected *Cdr1as* in formalin-fixed paraffin-embedded (FFPE) sections and directly visualized the therapeutic efficacy of AAV-delivered double-stranded circRNAs (ds-cRNAs)^32^ by reduced astrogliosis near transduced cells.

## Results

### Single molecule circRNA detection by RCA

To achieve circRNA-specific detection, we designed two partially complementary DNA probes: a padlock probe targeting the upstream back-splicing acceptor site and a primer probe complementary to the downstream back-splicing donor site (Fig.1a). Individual padlock and primer probes each forms 22-bp hybridization with the circRNA sequence of interest. When bound adjacently at the BSJ site, such a probe forms a nicked double-stranded DNA structure. The nick is sealed by T4 DNA ligase, circularizing the padlock probe to serve as a template for RCA (Fig. 1a). RCA is initiated from the primer probe using φ29 DNA polymerase, generating an amplicon harboring tandem repeats hybridized to the target circRNA. Fluorescent probes are then hybridized to the amplicon for imaging (Fig. 1a). Theoretically, although probes may hybridize to linear transcripts containing the same exonic sequences, the subsequent ligation and amplification are prevented due to steric constraints and the spatial separation of probe-binding sites on the linear RNA molecule (Fig. 1a).

**Fig. 1.**
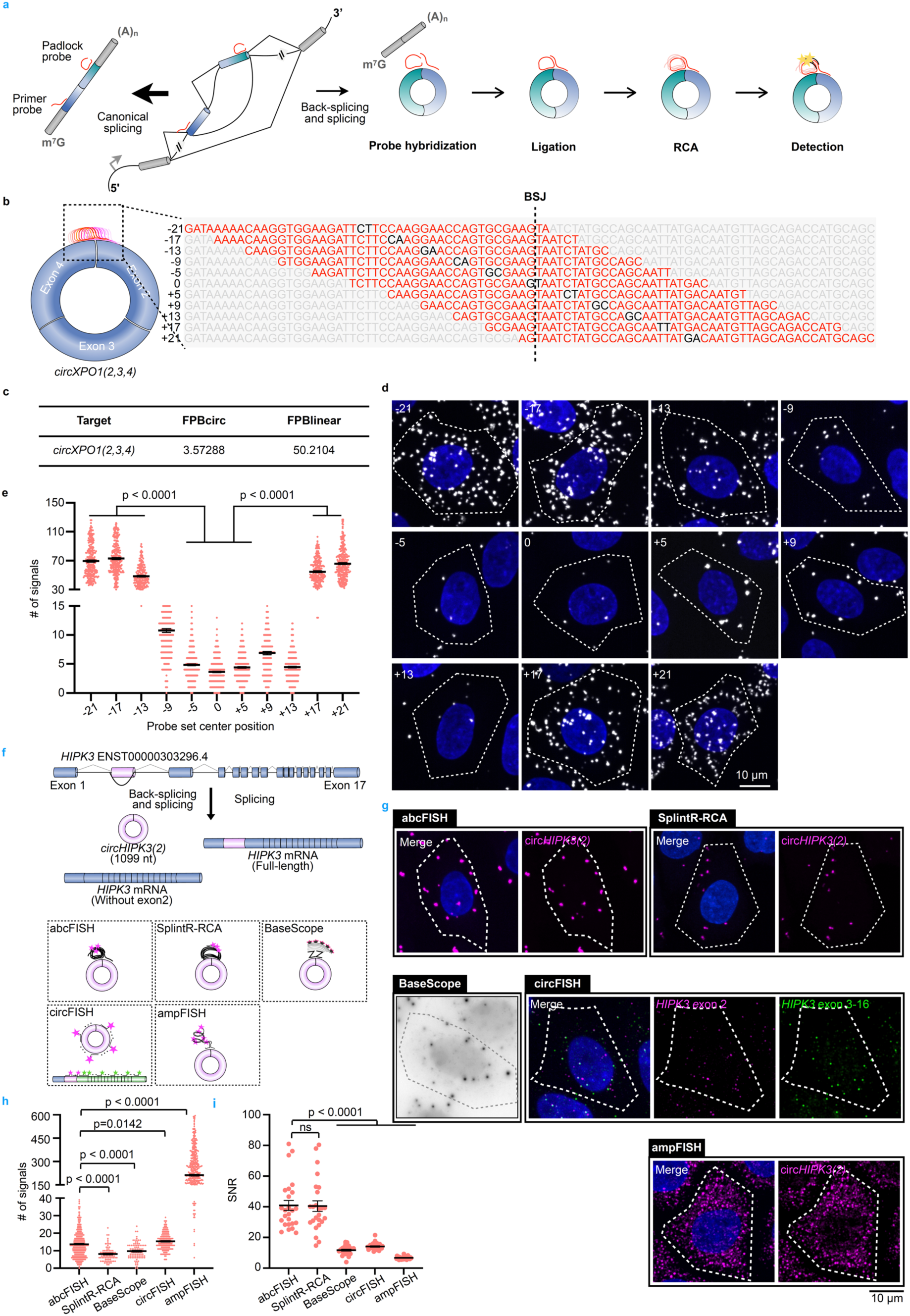
RCA enables single-molecule imaging of circRNAs. **a,** Schematic of the abcFISH for circRNA detection. A primer probe and a 5’ phosphorylated padlock probe bind adjacently across the BSJ, followed by ligation mediated by T4 DNA ligase, RCA amplification mediated by phi29 DNA polymerase, and detection by fluorescent probes for circRNA visualization. **b,** Schematic for systematic tiling of padlock and primer probe binding sites across the BSJ region of *circXPO1(2,3,4)* to determine the optimal hybridization position. The BSJ is indicated by a vertical dashed line at position 0. Binding sequences targeted by the padlock and primer probes are highlighted in red, with a 2-nucleotide gap shown in black between adjacent probe-binding regions. The central nucleotide position of each probe set is labeled on the left. **c,** Sequence abundance of *circXPO1(2,3,4)* and *XPO1* mRNA, showing FPB values for circRNA (FPB_circ_ = 3.57) and its linear counterpart (FPB_linear_ = 50.21). **d,** Representative abcFISH images of probe tiling around the BSJ shown in **b**, in HeLa cells. Cellular boundaries are indicated with white dashed lines. **e,** Statistics of abcFISH signals corresponding to probe positions shown in **d**. Data represent mean ± s.e.m.; n = 250 (-21), 271 (-17), 184 (-13), 229 (-9), 338 (-5), 314 (0), 317 (+5), 195 (+9), 359 (+13), 236 (+17), and 275 (+21) cells; *P* values were calculated using a one-way analysis of variance (ANOVA). **f,** Schematic of linear and circular *HIPK3* transcripts and detecting circRNAs using abcFISH, SplintR-RCA, BaseScope, circFISH and ampFISH. **g,** Representative images of *circHIPK3(2)* in cells using abcFISH, SplintR-RCA, BaseScope, circFISH and ampFISH. abcFISH, SplintR-RCA, circFISH, and ampFISH, employed fluorescent detection, with abcFISH and ampFISH utilizing Cy3-labeled probes, and circFISH probes tailed with Gold550 dUTP-labeled probes. BaseScope employed Fast red for brightfield detection. **h,** Statistics of *circHIPK3(2)* molecules per cell using abcFISH, SplintR-RCA, BaseScope, circFISH and ampFISH. abcFISH and circFISH detected comparable numbers of *circHIPK3(2)*, whereas SplintR-RCA and BaseScope detected fewer signals and ampFISH yielded non-specific signals. Data represent mean ± s.e.m.; n = 414 (abcFISH), 81 (SplintR-RCA), 92 (BaseScope), 181 (circFISH) and 529 (ampFISH) cells; *P* values were calculated using a one-way analysis of variance (ANOVA). **i,** Statistics of *circHIPK3(2)* SNR using abcFISH, SplintR-RCA, BaseScope, circFISH and ampFISH. abcFISH and SplintR-RCA achieved the highest SNR. Data represent mean ± s.e.m.; n = 25 cells per group; *P* values were calculated using a one-way analysis of variance (ANOVA).

To examine abcFISH specificity for circRNA imaging, we designed padlock probes tiling the BSJ of *circXPO1(2,3,4)*, a low-abundance circRNA (FPB_circ_ = 3.57) with expression markedly lower than its linear isoform (FPB_linear_ = 50.21) (Fig. 1b, c). FPB (fragments per billion bases) provides a base-normalized unit enabling direct comparison of circRNA and linear RNA expression in RNA-seq datasets^33^. Probes binding > 9 nucleotides (nt) from the BSJ center produced high non-specific signals, consistent with the off-target detection of abundant linear transcripts containing the same exon sequences (Fig. 1d, e). Notably, although probes spanning –9 to +9 nt yielded signals that were orders of magnitude lower than those binding farther from the BSJ, the –5 to +5 nt probes produced even more consistent signal numbers, indicating that this narrower region represents a specific hybridization window for circRNA detection (Fig. 1d, e, Extended Data Fig. 1g).

RNAfold predicted stem-loop structures flanking the BSJ site of *circXPO1(2,3,4)* (Extended Data Fig. 1a); however, these structures were unlikely to affect the detection efficiency of designed abcFISH probes (Fig. 1d, e). Analysis of published eCLIP-seq data^34^ revealed that several RBPs bound to *circXPO1(2,3,4)* with reads spanning the BSJ (Extended Data Fig. 1b, c), but with much lower abundance on *circXPO1(2,3,4)* than that on its linear RNA (Extended Data Fig. 1c), suggesting the high probe accessibility for this circRNA.

Next, we employed the established smFISH-based circFISH approach^16^ to validate abcFISH signals (Fig. 1d, e). circFISH^16^ used Gold550-labeled probes targeting the circularized exons of *circXPO1(2,3,4)*, and Red650-labeled probes specific to exons found only in linear transcripts (Extended Data Fig. 1d). Circular transcripts were quantified by counting Gold550-only puncta (Extended Data Fig. 1e, f). Of note, the short length of *circXPO1(2,3,4)* (307 nt) allowed for only 14 probes, fewer than the > 20 probes typically required for robust smFISH detection, thus resulting in a decreased SNR and non-specific signals (Extended Data Fig. 1e) compared to abcFISH (Fig. 1d). Furthermore, circFISH probes also detected nascent *XPO1* pre-mRNAs, visible as bright nuclear foci (Extended Data Fig. 1e), which challenged the discrimination of true circRNA signals. Despite these limitations, circFISH quantification of Gold550-only puncta yielded an average of approximately 4 copies of *circXPO1(2,3,4)* per cell (Extended Data Fig. 1e-g). This result aligns with the copy number quantified by abcFISH using probes spanning –5 to +5 BSJ region (Fig. 1e, Extended Data Fig. 1g) and correlates with the RNA-seq FPB_circ_ value (Fig. 1c), demonstrating 28 nucleotides to the right or left of the BSJ site are targetable by abcFISH probes (Extended Data Fig. 1h). These analyses together show that abcFISH likely offers superior specificity and a simpler design for imaging circRNAs compared to the smFISH-based circFISH. Of note, abcFISH probes (except those used in Fig. 1b) used in this study were centered on BSJ.

### RCA-based abcFISH outperforms existing methods for circRNA imaging

Next, we performed a head-to-head comparison of abcFISH against existing circRNA imaging methods: the smFISH-based circFISH^16^ as mentioned above, the hybridization chain reaction (HCR)-based amplification FISH (ampFISH) ^28, 35^, another RCA-based method (herein referred to as SplintR-RCA) ^27^, and commercial BaseScope^29^. ampFISH is an enzyme-free method based on HCR for signal amplification with minimal background^35, 36^, and has been used to image cellular and viral circRNAs^28^. SplintR-RCA directly hybridizes padlock probes on the circRNA BSJ with the ligation step mediated by SplintR ligase^27^. BaseScope detects circRNAs through alkaline phosphatase after multiple amplification steps^29^ (Fig. 1f). We compared these techniques by targeting the cancer-associated *circHIPK3(2)*^37^ in HeLa cells (Fig. 1f).

All methods produced cytoplasmic signals (Fig. 1g). Quantitation showed that abcFISH and circFISH detected similar number of *circHIPK3(2)* molecules per cell, demonstrating strong concordance and confirming abcFISH’s sensitivity. SplintR-RCA and BaseScope detected fewer *circHIPK3(2)* than abcFISH, likely reflecting limited ligation efficiency when the substrate carries a 5′ phosphorylated dC nucleotide^38^ for SplintR-RCA and decreased efficiency brought by multiple-step chemistry for BaseScope. In contrast, ampFISH generated hundreds of non-specific signals per cell (Fig. 1g, h) as previously reported^28^, indicating substantial off-target amplification and a high false-positive rate. Moreover, both abcFISH and SplintR-RCA achieved the highest SNR, a direct benefit of RCA (Fig. 1i).

To demonstrate the specificity of the direct circRNA imaging methods, we knocked down *circHIPK3(2)* expression prior to imaging (Extended Data Fig. 2a). *circHIPK3(2)* expression level decreased with circRNA-specific siRNA transfection, as measured by qPCR (Extended Data Fig. 2b). abcFISH, SplintR-RCA and BaseScope all showed high specificity, as the signals dropped according to decreased *circHIPK3(2)* expression (Extended Data Fig. 2c, d). In contrast, ampFISH produced hundreds of signals per cell regardless of siRNA treatment (Extended Data Fig. 2c, d), indicating that the vast majority of ampFISH signals was indeed non-specific, and thus this HCR-based method is not suitable for circRNA imaging.

Critically, abcFISH overcomes key limitations of existing techniques. Unlike circFISH, which requires circRNAs to be long enough (> ∼300 nt) to accommodate multiple probes, abcFISH imposes no size constraints as it targets only the short sequence spanning BSJ. Additionally, the RCA-amplified products contain universal sequences within the padlock probe backbone, allowing the same fluorescent detection probes to be reused for different circRNA targets; only the BSJ-binding sequence for the padlock needs customization, showing flexibility in probe design for different circRNA targets. Unlike SplintR-RCA, abcFISH applies T4 DNA ligase instead of SplintR ligase, lowering the cost and preventing ligated substrate constraints. Furthermore, with shorter padlock probes than those used in SplintR-RCA, abcFISH costs only approximately 28 USD per reaction. Although BaseScope provides mature circRNA imaging solution, the brightfield imaging nature limits its SNR and imaging multiplexity, and it costs higher than the other listed methods (Extended Data Fig. 3a). The RCA system, combined with the direct BSJ targeting, makes abcFISH highly convenient, low-cost and specific, eliminating the complex computational post-processing required for signal discrimination in circFISH.

### abcFISH provides quantitative accuracy across diverse circRNAs

To further evaluate the quantitative accuracy of abcFISH, we selected four additional circRNAs, *circSPECC1(4)*, *circASPH(2,3)*, *circCDYL(2)* and *circPOLR2A(9,10)*, with various expression levels in HeLa cells (Extended Data Fig. 4a). We performed parallel imaging of *circCDYL(2*) and *circSPECC1(4)* using abcFISH and circFISH (Extended Data Fig. 4b-f). The other two targets, *circASPH(2,3)* and *circPOLR2A(9,10)*, are only 219 nt and 336 nt long, respectively, rendering them too short for reliable detection by circFISH. We also used qPCR to measure the absolute circRNA abundance from bulk RNA. Intact circularized exons were cloned into plasmids with the BSJ positioned centrally to generate standard curves from serial dilutions of these plasmids (Extended Data Fig. 4g). The copy number of circRNA per cell was calculated based on the standard curves and input cells (Extended Data Fig. 4h).

Both abcFISH and circFISH detected cytoplasmic signals for their respective targets. abcFISH successfully detected the low-abundance *circCDYL(2)* and *circPOLR2A(9,10)*, quantifying an average ∼ 5 copies per cell, which underscores its high sensitivity (Extended Data Fig. 4b,c). For targets measurable by both techniques (*circCDYL(2*) and *circSPECC1(4)*), abcFISH showed comparable but slightly higher signals than circFISH, confirming it as a robust quantitative imaging approach (Extended Data Fig. 4i). Of note, the copy numbers of these examined circRNAs were also confirmed with qPCR, showing a comparable level (Extended Data Fig. 4i).

Collectively, the consistent quantitative performance of abcFISH across circRNAs of varying abundances and lengths, combined with its unique ability to preserve spatial context, establishes it as a robust and specific approach for circRNA quantification that avoids the amplification artifacts and template biases inherent to PCR-based methods.

### Multiplexed and quantitative circRNA imaging across cell types

Since abcFISH showed high accuracy in quantifying circRNAs and overcame the size limitation of the canonical imaging method (Extended Data Fig. 4), we next investigated whether abcFISH enables the simultaneous detection of multiple circRNAs within the same cells. We designed three spectrally distinct fluorophores (AF488, AF568 and AF647) targeting *circSPECC1(4)*, *circASPH(2,3)* and *circCDYL(2*), respectively, and distinct cytoplasmic signals were shown for each circRNA, simultaneously (Fig. 2a). The three circRNAs showed expected abundance (Fig. 2b), aligning with their FPB and qPCR values (Extended Data Fig. 4a, h).

**Fig. 2.**
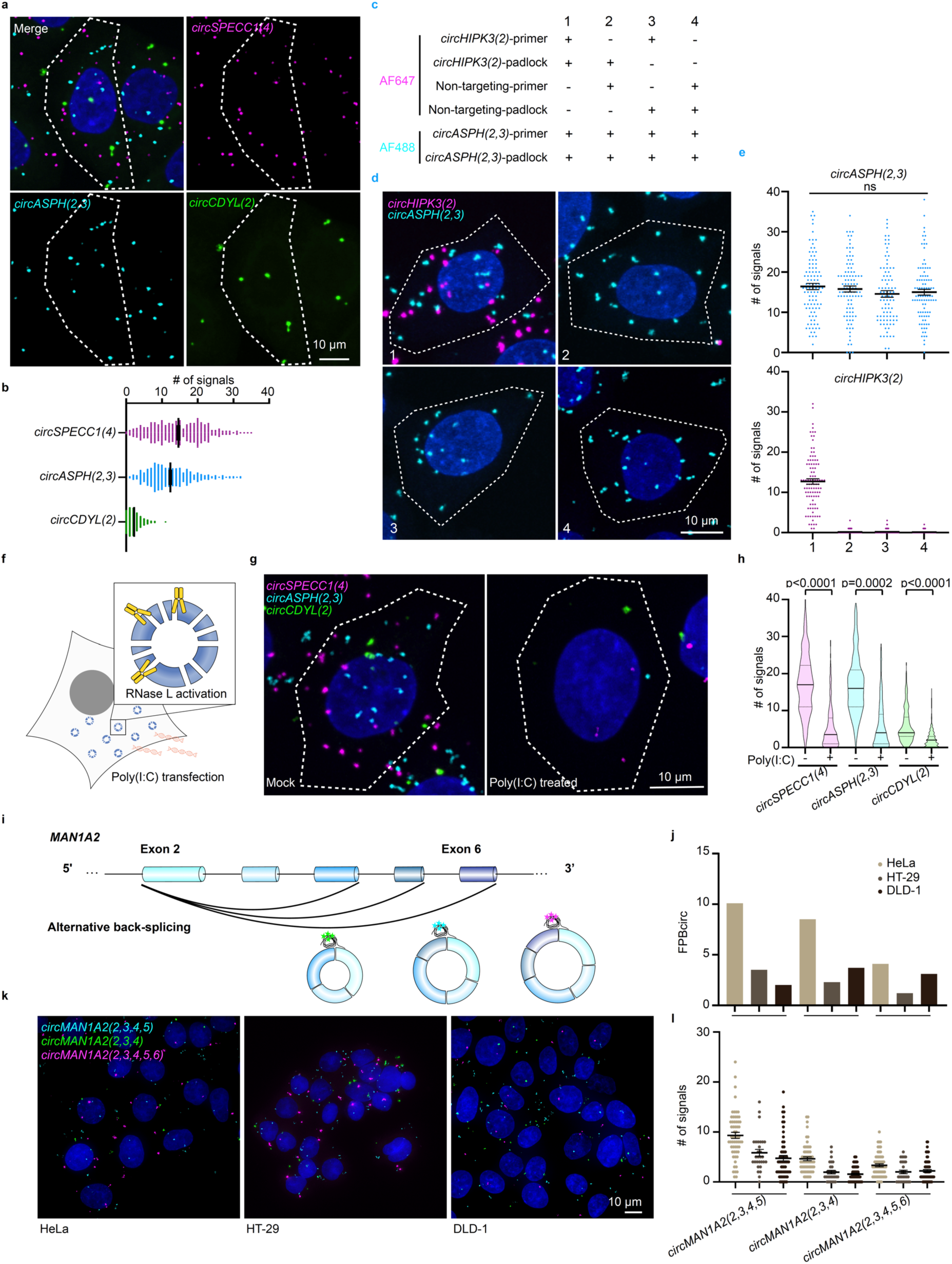
abcFISH enables detection of circRNA degradation and alternative circularization. **a,** Representative images of *circSPECC1(4)*, *circASPH(2,3)* and *circCDYL(2*) simultaneously labeled by abcFISH in HeLa cells using AF647, AF568 and AF488, respectively. Magenta, *circSPECC1(4)* signals. Cyan, *circASPH(2,3)* signals. Green, *circCDYL(2)* signals. **b,** Statistics of single-cell circRNA copy numbers showed distinct expression levels of *circSPECC1(4)*, *circASPH(2,3)* and *circCDYL(2)*. Data represent mean ± s.e.m; n = 318 cells per group. **c,** Simultaneous imaging of *circASPH(2,3)* with on-targeting probes and *circHIPK3(2)* under 4 conditions (1. both on-targeting, 2. primer non-targeting, 3. padlock non-targeting, 4. both non-targeting). **d,** Representative images of *circASPH(2,3)* (cyan, AF488 channel) and *circHIPK3(2)* (magenta, AF647 channel) in HeLa cells using on-targeting or non-targeting abcFISH probes for *circHIPK3(2)*. **e,** Statistics of detected *circASPH(2,3)* and *circHIPK3(2)* signals per cell shown in **d**. *circASPH(2,3)* signals were not affected by non-targeting probes against *circHIPK3(2)*, whereas *circHIPK3(2)* signals were almost completely abolished when either the padlock or the primer was non-targeting, and fully absent when both were non-targeting. Data represent mean ± s.e.m.; n = 98 (1), 98 (2), 88 (3) and 99 (4) cells;; *P* values were calculated using a one-way analysis of variance (ANOVA). **f,** Schematic of innate immune activation by poly(I:C). Upon treatment of poly(I:C), cellular endonuclease RNase L is activated, which leads to cleavage of circRNAs. The global circRNA level decreases substantially due to its slow biogenesis. **g,** Representative abcFISH images of *circSPECC1(4)*, *circASPH(2,3)* and *circCDYL(2*) in HeLa cells before and 4 hours after 1 µg/mL poly(I:C) transfection, showing RNase L–mediated circRNA degradation. **h,** Statistics of circRNA signals per cell confirmed rapid circRNA loss upon immune stimuli in HeLa cells. Data represent mean ± s.e.m; n = 290 (mock) and 402 (poly(I:C) treated) cells; *P* values were calculated using two-tailed Student’s t-test. **i,** Schematic of alternative back-splicing at the MAN1A2 locus. Three highly expressed circular isoforms were alternative circularized with distinct exon compositions. Multiplex abcFISH with different fluorophores was performed to image *circMAN1A2(2,3,4)*, *circMAN1A2(2,3,4,5)* and *circMAN1A2(2,3,4,5,6)*. **j,** Expression of distinct *circMAN1A2* isoforms across HeLa, HT29, and DLD-1 cells based on previous ribo– RNA-seq data^40^. **k,** Representative abcFISH images of three *circMAN1A2* AC isoforms in HeLa, HT29 and DLD-1 cells, showing their cytoplasmic localization. Magenta, *circMAN1A2(2,3,4,5,6)* signals. Cyan, *circMAN1A2(2,3,4,5)* signals. Green, *circMAN1A2(2,3,4)* signals **l,** Statistics of single-cell circRNA copy numbers for three *circMAN1A2* AC isoforms, showed concordance with RNA-seq–derived expression levels in **j**. Data represent mean ± s.e.m; n = 60 (HeLa), 29 (HT-29) and 61 (DLD-1) cells.

To further validate the specificity of abcFISH in multiplex circRNA imaging, we simultaneously imaged *circHIPK3(2)* and *circASPH(2,3)*. For *circASPH(2,3)*, we used on-targeting probes as controls, whereas for *circHIPK3(2)* we replaced either the padlock or the primer probe with non-targeting sequences (Fig. 2c). Note that the use of non-targeting padlock or primer probes specifically eliminated *circHIPK3(2)* signals while leaving *circASPH(2,3)* detection accurate in the other channel (Fig. 2d, e), confirming that abcFISH specifically images circRNAs only with on-targeting probes and that this specificity is maintained during multiplex imaging.

We next applied imaging to visualize the fate of these circRNAs. We treated cells with poly(I:C) to activate the endonuclease RNase L, which is known to globally degrade circRNAs^10^, followed by multiplexed imaging of circRNAs (Fig. 2f). As expected, after 4 hours of poly(I:C) transfection, we observed a remarkable reduction in signals for all three circRNAs (Fig. 2g, h).

Alternative circularization (AC) allows a single gene locus to produce multiple circRNA isoforms with distinct exon compositions, a phenomenon observed for >50% of circRNA-producing loci^17, 39^. We applied abcFISH to directly visualize AC by simultaneously imaging three isoforms of *circMAN1A2*: *circMAN1A2(2,3,4)*, *circMAN1A2(2,3,4,5)* and *circMAN1A2(2,3,4,5,6)*, which are associated with colon cancer^40^ (Fig. 2i, j). Isoform-specific abcFISH successfully resolved these variants in three different cell lines, HeLa, HT29 and DLD-1 cells, with cytoplasmic localization patterns (Fig. 2k). The quantified signal counts for each isoform (Fig. 2l) correlated with their respective expression levels as determined by ribosomal RNA-depleted RNA sequencing (ribo–RNA-seq) across cell types^40^ (Fig. 2j). These results further support the capability of abcFISH for multiplexed, quantitative, and isoform-specific visualization of circRNAs. Collectively, abcFISH signal numbers of these circRNAs in HeLa cells were positively correlated with their FPB_circ_ values, further demonstrating the accuracy of abcFISH (Extended Data Fig. 4j).

### abcFISH enables visualization and quantification of *circPOLR2A(9,10) in vivo*

To explore the utility of abcFISH for circRNA imaging *in vivo*, we visualized *circPOLR2A(9,10)* in various tissues from a KI mouse model. In this model, a human *circPOLR2A(9,10)* expression cassette was inserted into the *Rosa26* locus under the control of the EF1α promoter, producing both *circPOLR2A(9,10)* and the full-length *EGFP* mRNA (Fig. 3a). As *circPOLR2A(9,10)* is not expressed in wild-type (WT) mice, abcFISH signals in KI mice brain sections were highly specific, without background detected in WT controls (Fig. 3b). Given the higher tissue autofluorescence than that in cultured cells^41^, we compared three fluorophores (AF488, AF568, AF647) and found AF647 yielded the highest SNR due to the minimal tissue autofluorescence in the far-red spectrum (Fig. 3c, d). This enables an optimized protocol for *in vivo* imaging.

**Fig. 3.**
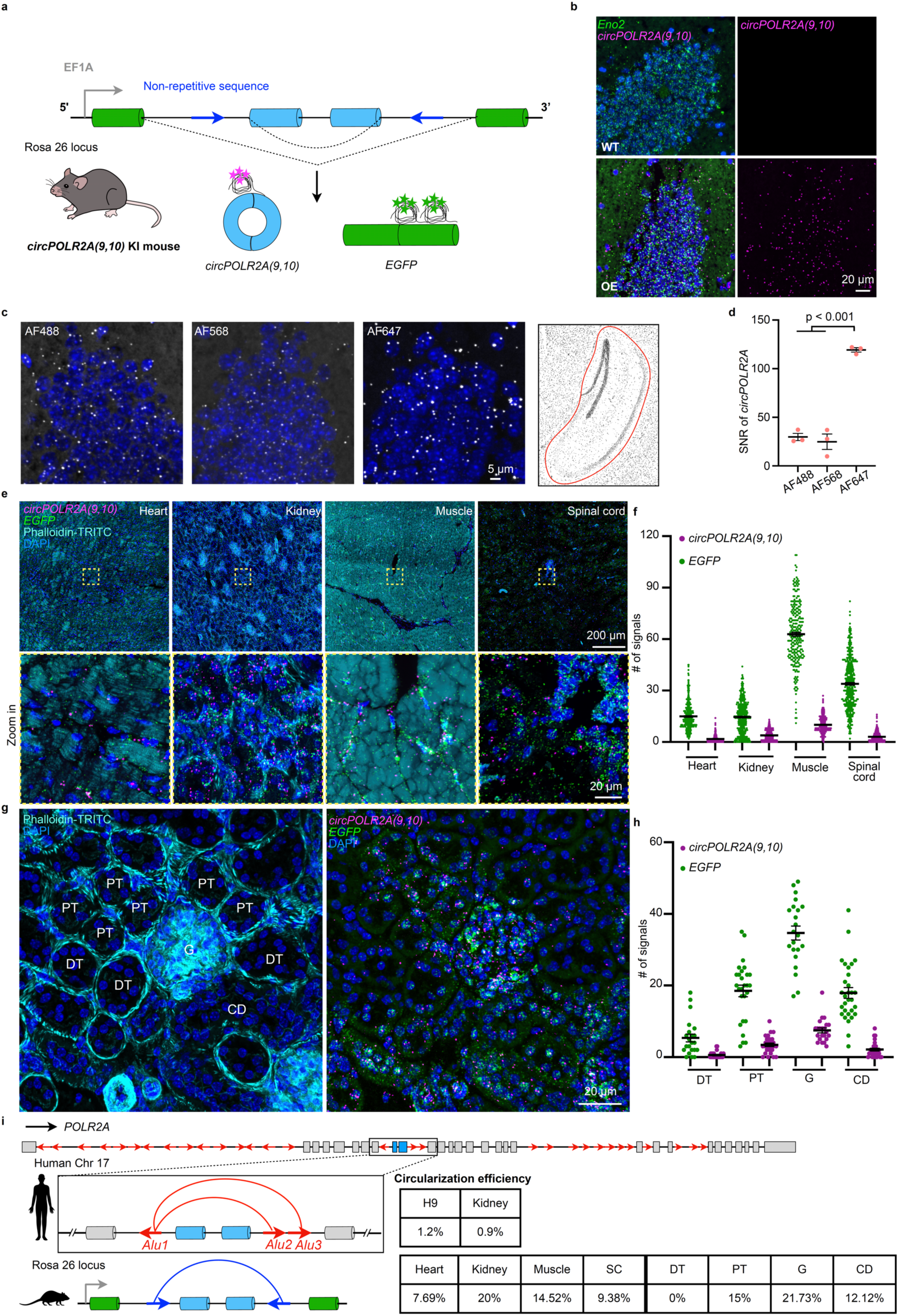
*In vivo* abcFISH reveals heterogeneous expression of *circPOLR2A(9,10)* across tissues. **a,** Schematic of the human *circPOLR2A(9,10)* KI mouse model. An expression cassette containing human *POLR2A* exons 9 and 10 flanked by non-repetitive sequences (blue arrows) was inserted into the Rosa26 locus under the control of an EF1α promoter, enabling simultaneous production of *circPOLR2A(9,10)* and full-length *EGFP* mRNA. The generated *circPOLR2A(9,10)* was detected via abcFISH targeting BSJ, while *EGFP* mRNA was detected via abcFISH targeting two sites within downstream half *EGFP* exon in the following experiment. **b,** Representative images of specific abcFISH detection of *circPOLR2A(9,10)* in mouse brain sections. Neurons in the granule cell layer of the dentate gyrus (GrDG) were marked by *Eno2* mRNA. No signal was observed in WT mice (3-month-old mice, 3M), demonstrating high specificity of abcFISH *in vivo*. Green, *Eno2* mRNA signals. Magenta, *circPOLR2A(9,10)* signals. **c,** Representative abcFISH images of fluorophore optimization for circRNA imaging *in vivo*. *circPOLR2A(9,10)* in KI-mouse brain sections was targeted with the same primer and padlock probes, but hybridized with detection probes labeled with AF488, AF568 and AF647. The images showed partial GrDG shown in blue square on the right panel. White, *circPOLR2A(9,10)* signals. **d,** Statistics of *circPOLR2A(9,10)* SNR using different fluorophores shown in **c.** AF647 detection probes yielded the highest SNR. SNR was calculated within the hippocampal region of interest (ROI) outlined in red in the right panel of **c**. Each data point represents the mean SNR from two hippocampal regions per brain section. Data represent mean ± s.e.m.; n = 3 brain sections per group; *P* values were calculated using two-tailed Student’s t-test. **e,** Representative abcFISH images of *circPOLR2A(9,10)* and *EGFP* mRNA across multiple tissues—heart, kidney, muscle, and spinal cord—from KI mice. Cell boundaries were visualized by phalloidin staining. ROIs indicated by yellow dashed squares, are shown at higher magnification in the bottom panels. Green, *EGFP* mRNA signals. Magenta, *circPOLR2A(9,10)* signals. Cyan, phalloidin staining. **f,** Statistics of *circPOLR2A(9,10)* (purple dots) and *EGFP* mRNA (green dots) copy number per cell across tissues shown in **e**. Data represent mean ± s.e.m.; n = 361 (heart), 411 (kidney), 259 (muscle) and 419 (spinal cord) cells. **g,** Representative abcFISH images of *circPOLR2A(9,10)* and *EGFP* mRNA expression in KI-mouse kidney. Glomerulus (G), proximal tubules (PT) and collecting ducts (CD) were identified by phalloidin staining, but not in distal tubules (DT). Green, *EGFP* mRNA signals. Magenta, *circPOLR2A(9,10)* signals. Cyan, phalloidin staining. **h,** Statistics of *circPOLR2A(9,10)* (purple dots) and *EGFP* mRNA (green dots) copy number per cell in KI-mouse kidney shown in **g**. Data represent mean ± s.e.m.; n = 22 (DT), 27 (PT), 22 (G) and 30 (CD) cells. **i,** Circularization efficiency of *circPOLR2A(9,10)* in H9 cell line, human kidney samples, KI-mouse tissues and kidney substructures. The left panel shows the different intronic pairing of POLR2A in human genome which is flanked by multiple *Alu* elements (indicated by red arrows) and in KI mouse Rosa 26 locus. Human data are derived from CIRCpedia v3^69^ (the top table). Mouse data are derived from tissue-level quantifications in **f** (first four columns) and kidney substructure quantifications in **h** (last four columns) (the bottom table). G, glomerulus; PT, proximal tubule; CD, collecting duct; DT, distal tubule.

We next performed abcFISH on KI mouse tissue sections from heart, kidney, muscle and spinal cord, using phalloidin to outline tissue architectures. Meanwhile, the co-expressed *EGFP* mRNA served as an intrinsic reporter for transcription of the KI cassette (Fig. 3a), allowing simultaneous single-molecule detection of both *circPOLR2A(9,10)* and *EGFP* mRNA within tissue sections (Fig. 3e). As expected, this engineered cassette was expressed at different levels across tissues as quantified by the *EGFP* mRNA, as higher transcription of the EF1α promoter led to higher accumulation of *EGFP* mRNA and *circPOLR2A(9,10)* (Fig. 3f). Notably, the circularization efficiency, calculated as the median ratio of *circPOLR2A(9,10)* to *EGFP* mRNA, observed in this mouse model (ranging from 7.69% in heart to 20% in kidney; Fig. 3i) was substantially higher than that of most endogenous circRNAs^42^ (typically <1%) and *circPOLR2A(9,10)* in human cells (Fig. 3i). This elevated back-splicing efficiency could be largely due to the application of a human intronic pairing in this KI cassette without an endogenous sequence competition in the mouse genome, thus providing a strong *in vivo* support for the previous view that the alternative formation and competition of inverted repeated (mostly *Alu*) pairs are key drivers of alternative circularization (Fig. 3i)^17, 39^.

Nonetheless, across the tissues, the circularization varied, and within the kidney, *circPOLR2A(9,10)* exhibited distinct expression patterns even among anatomical substructures. It was differently expressed in glomerulus (G), proximal tubules (PT) and collecting ducts (CD), but absent in distal tubules (DT) (Fig. 3g, h). These structures were identified by phalloidin-stained cytoskeleton^43^ (Fig. 3g), revealing a heterogeneous landscape of circRNA expression at the tissue level. This heterogeneity likely reflects the multilayer regulation in circRNA biogenesis, including transcription rate, RNA turnover and cell division rate across these structures^42, 44^.

### Heterogeneity of *circPOLR2A(9,10)* expression across hippocampal cell types

Although circRNAs were thought to be highly enriched in the mammalian brain^11, 19^, to what extent their expression varies across diverse brain cell types has remained unclear due to the lack of robust detection tools in tissues. Combining abcFISH with a molecular spatial atlas of the central nervous system^45^, we studied *circPOLR2A(9,10)* expression across cell types in the KI-mouse brain.

We focused on major brain cell types, including neurons, astrocytes, oligodendrocytes, microglia, vascular endothelial cells, vascular smooth muscle cells, pericytes and choroid plexus epithelial (CPE) cells (Fig. 4a), annotated with representative molecular markers (Extended Data Fig. 5a). We observed remarkable heterogeneity of *circPOLR2A(9,10)* across cell types. For example, CPE cells (marked by *Ttr* mRNA) exhibited the highest *circPOLR2A(9,10)* levels while oligodendrocytes (marked by *Opalin* and *Mal* mRNAs) showed very low expression (Fig. 4b, c). Although their expression was generally coordinated with the expression of this KI cassette (Fig. 3a), the circularization efficiency across these cells varied as we observed across tissues (Extended Data Fig. 5a, Fig. 3i).

**Fig. 4.**
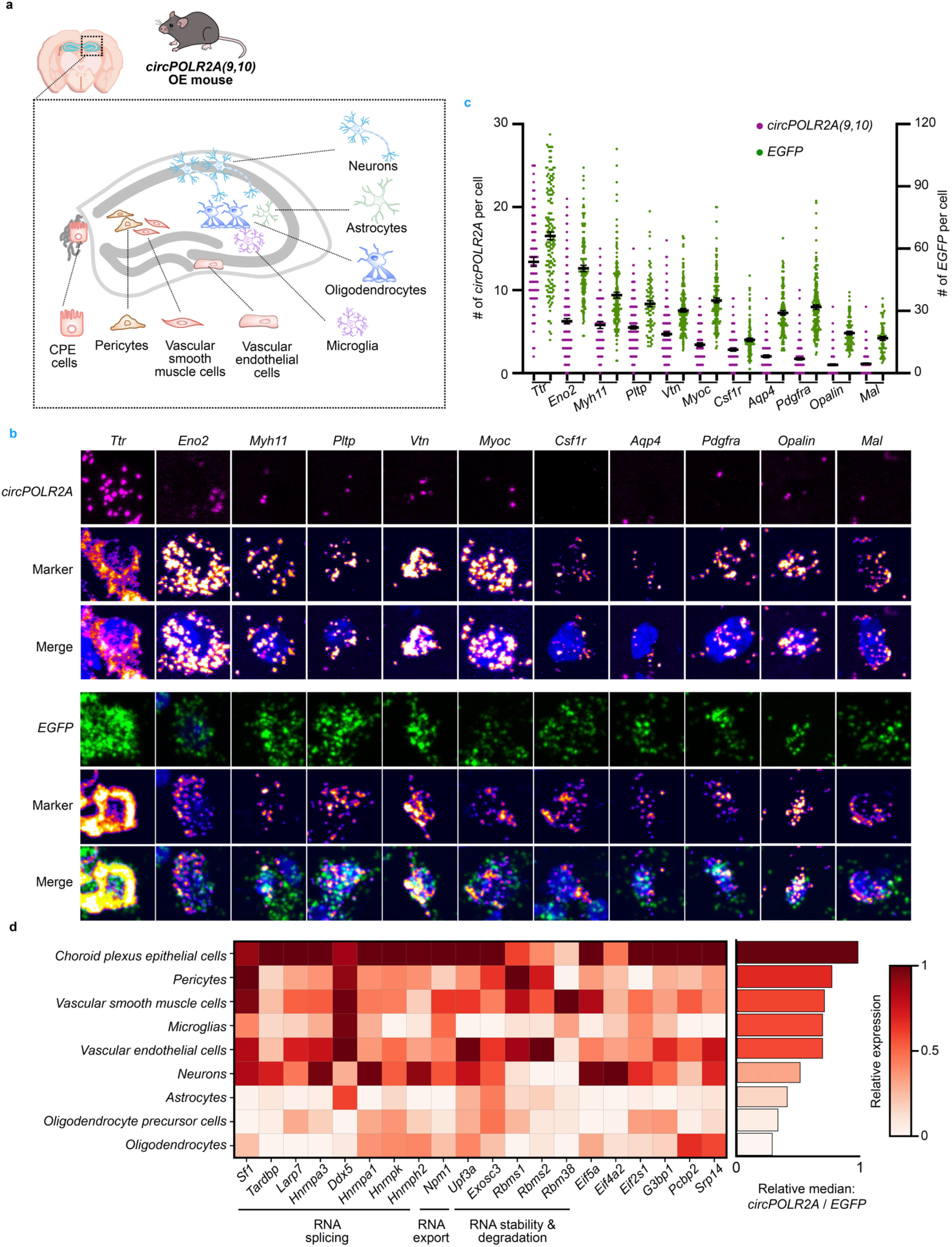
*CircPOLR2A(9,10)* expression is regulated by potential combinatorial effects of RBPs. **a,** Schematic of cell type-specific mapping of *circPOLR2A(9,10)* expression in the KI mouse hippocampus. Neurons, astrocytes, oligodendrocytes, microglia, vascular endothelial cells, vascular smooth muscle cells, pericytes and CPE cells localized across hippocampus were included. **b,** Representative abcFISH images of *circPOLR2A(9,10)* and *EGFP* mRNA expression in cell types shown in **a**. *circPOLR2A(9,10)* (top panels) and *EGFP* mRNA (bottom panels) were separately imaged with AF647 probes with marker RNAs detected with either AF568 or AF647 probes for cell type identification. Green, *EGFP* mRNA signals. Magenta, *circPOLR2A(9,10)* signals. Flame, marker RNA signals. **c,** Statistics of *circPOLR2A(9,10)* (purple dots, left y-axis) and *EGFP* mRNA (green dots, right y-axis) copy numbers per cell type shown in **b**. Data represent mean ± s.e.m.; for circRNA quantification, n = 85 (*Ttr*), 211 (*Eno2*), 64 (*Myh11*), 222 (*Pltp*), 178 (*Vtn*), 86 (*Myoc*), 140 (*Csf1r*), 109 (*Aqp4*), 182 (*Pdgfra*), 153 (*Opalin*), 181 (*Mal*) cells; for *EGFP* mRNA quantification, n = 159 (*Ttr*), 122 (*Eno2*), 146 (*Myh11*), 81 (*Pltp*), 176 (*Vtn*), 125 (*Myoc*), 100 (*Csf1r*), 118 (*Aqp4*), 173 (*Pdgfra*), 72 (*Opalin*), 62 (*Mal*) cells **d,** Heatmap of the top 20 RBPs most strongly correlated with the *circPOLR2A(9,10)*-to-*EGFP* expression ratio across cell types. These RBPs were analyzed from previously screened 103 RBPs^46^, with Gene Ontology (GO) annotated functions are provided below.

We next investigated factors likely contributing to this heterogeneous expression. Given that RNA binding proteins (RBPs), known modulators of back-splicing^46–48^, are highly variable across brain cells^49^, we analyzed RBP expression using publicly available single-cell data from the Allen Brain Atlas^50^ (Extended Data Fig. 5a, b). We cross-referenced 103 RBPs previously identified in a loss-of-function screen for circRNA turnover^46^ and identified the top 20 whose expression was most highly correlated with circularization efficiencies across these cells (Fig. 4d). Among these, HNRNPK (heterogeneous nucleoprotein K) was recently found to bind flanking introns and enhance circRNA biogenesis^51^. RNA helicase DDX5 could unwind RNA structures near the BSJ facilitating back-splicing^52^. In addition, TARDBP, HNRNPA3, and LARP7 showed enrichment in the proximity labeling assay enabled by TAPRIP, among the candidates binding to the plasmid encoding circmCherry^51^. Cell types exhibiting high *circPOLR2A(9,10)* expression, such as CPE cells, showed elevated expression levels of these proteins (Fig. 4d, Extended Data Fig. 5b), implying that RBP heterogenous expression may contribute to the observed circRNA heterogeneity across brain cell types. Notably, no single RBP showed a linear correlation with *circPOLR2A(9,10)* levels, supporting a model of combinatorial control of circRNA by multiple RBPs *in vivo*. This finding is consistent with previous work in *Drosophila*, where the Laccase2 circRNA level was regulated by multiple hnRNP and SR (serine-arginine) proteins ^53^.

### Multiplexed circRNA imaging revealed subregional enrichment in the hippocampus

We next applied the multiplexed abcFISH to map spatial distribution of endogenous circRNAs in the mouse hippocampus, simultaneously. These include *circZfp148(2,3,4)*, *circAnks1b(5,6,7,8)*, and the well-studied *Cdr1as*, which are expressed at different levels (Extended Data Fig. 6a, Fig. 5a). Each circRNA exhibited a distinct spatial distribution (Fig. 5a, Extended Data Fig. 6b). *circZfp148(2,3,4)* exhibited sparse and diffuse signals across the hippocampus, potentially reflecting low abundance. In contrast, both *circAnks1b(5,6,7,8)* and *Cdr1as* were enriched in the cornu ammonis (CA) regions and dentate gyrus granule cell layer (GCL), but displayed divergent localization patterns (Fig. 5a). Notably, *Cdr1as* was predominantly cytoplasmic, while >60% of *circAnks1b(5,6,7,8)* signals were nuclear (Fig. 5b, c), suggesting their distinct functions. Interestingly, their linear counterparts, *Anks1b* and *Cdr1os* (the mouse homolog of human *LINC00632*^15^), exhibited distinct localization patterns (Extended Data Fig. 7a-c), likely reflecting different mechanisms for linear and circular RNAs derived from the same gene loci. Furthermore, *Cdr1as* exhibited specific cytoplasmic patterns *in vivo*, and was highly concentrated in the CA1 stratum pyramidale but diffused in CA2 and CA3 (Fig. 5a, Extended Data Fig. 6b).

**Fig. 5.**
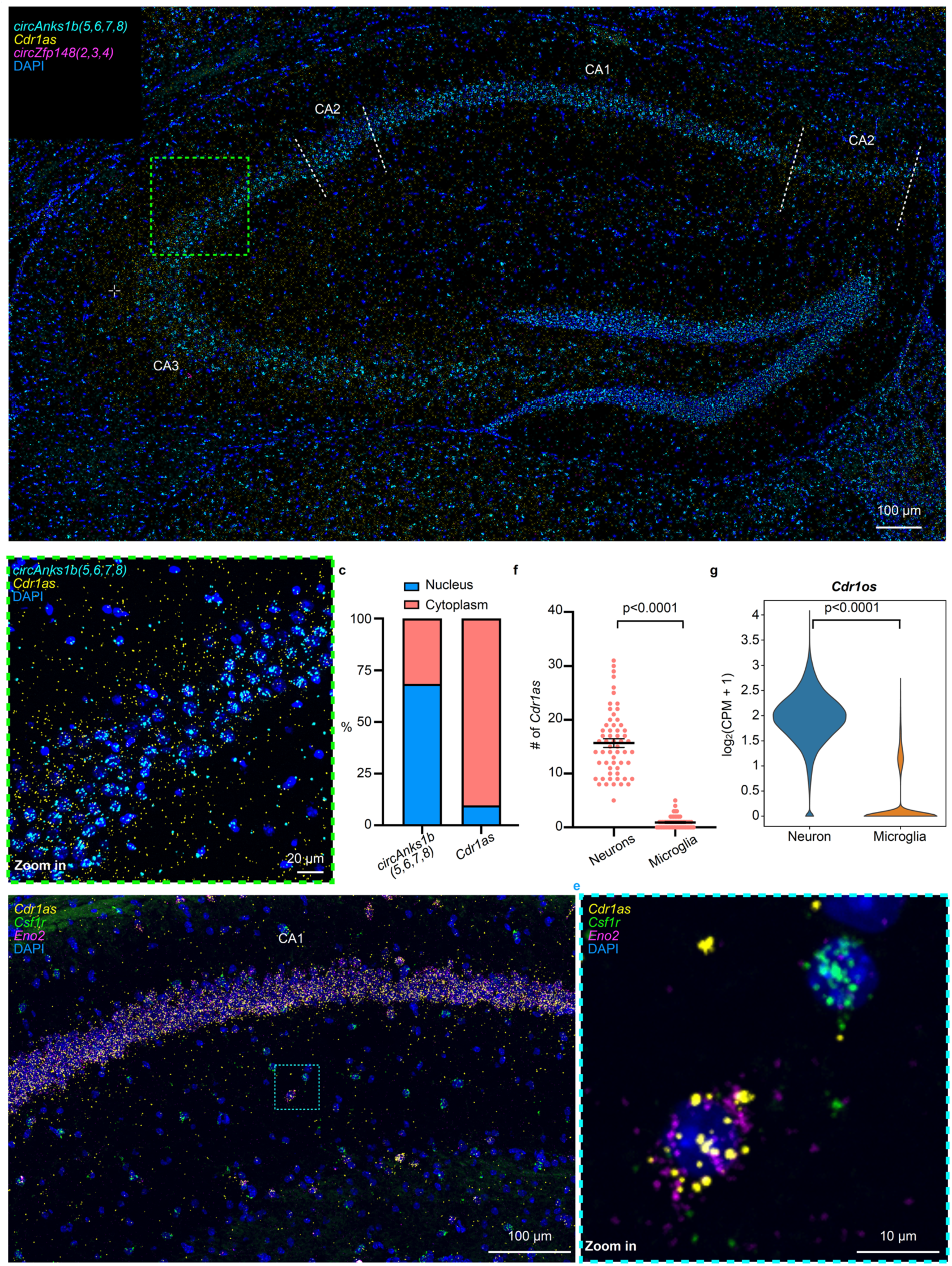
abcFISH reveals endogenous circRNA heterogeneity *in vivo*. **a,** The representative image of multiplex circRNA imaging in WT mouse (3-month-old) hippocampus. The ROI indicated by green dashed squares is shown at higher magnification in **b**. Subregions CA1, CA2, and CA3 are demarcated by white dashed lines. Yellow, *Cdr1as* signals. Cyan, *circAnks1b(5,6,7,8)* signals. Magenta, *circZfp148(2,3,4)* signals. **b,** Higher-magnification view of the ROI from **a**, revealing distinct subcellular localization patterns. *Cdr1as* displays a predominantly cytoplasmic distribution, while *circAnks1b(5,6,7,8)* exhibits significant nuclear enrichment. Yellow, *Cdr1as* signals. Cyan, *circAnks1b(5,6,7,8)* signals. **c,** The percentage of circRNA subcellular localization in the cytoplasm versus the nucleus. Data were analyzed from 60 cells shown in right panels of **b**. *Cdr1as* showed distribution in the cytoplasm while *circAnks1b(3,4,5,6,7,8)* exhibited dominant nuclear enrichment. **d,** The representative abcFISH image of detecting single circRNAs across cell types in WT mouse (3M) hippocampus. The ROI indicated by cyan dashed squares is shown at higher magnification in **e**. Neurons and microglia were marked by *Eno2* and *Csf1r* mRNA, respectively. Green, *Csf1r* mRNA signals. Magenta, *Eno2* signals. Yellow, *Cdr1as* signals. **e,** Higher-magnification view of the ROI from **d**, revealing preferential expression of *Cdr1as* in neurons than in microglia. **f,** Statistics of cell-type distribution of circRNAs shown in **e**. *Cdr1as* was more abundant in neurons than in microglia. Data represent mean ± s.e.m.; for *Cdr1as*, n = 59 (*Eno2*) and 57 (*Csf1r*) cells; *P* values were calculated using two-tailed Student’s t-test. **g,** Analysis of *Cdr1os* expression in microglia and neurons. The data were analyzed from Allen Brain Atlas. *P* values were calculated using Wilcoxon test.

To further evaluate cell-type-specificity, we co-imaged *Cdr1as* with cell markers (*Eno2* mRNA for neurons, and *Csf1r* mRNA for microglia; Fig. 5d), showing that *Cdr1as* was significantly more abundant in neurons than in microglia (Fig. 5e, f). We next investigated whether this heterogeneity is related to differential expression of its parental gene, *Cdr1os*, between these cell types. Expression analysis of *Cdr1os* confirmed its predominant expression in neurons (Fig. 5g), suggesting that *Cdr1as* biogenesis is linked to *Cdr1os* transcription.

### *Cdr1as* and *Cyrano* dominantly localize in neuronal soma with correlated expression

*Cyrano*, a lncRNA, promotes miR-7 degradation and facilitates *Cdr1as* accumulation, supported by knockout models and biochemical studies^31^. However, direct visualization of their spatial interaction at the single-molecule level in individual cells has been lacking. We applied abcFISH to investigate their spatial and quantitative relationship.

Methanol has been commonly used in spatial transcriptomics for sample permeabilization and preservation^25, 54^, but often results in soluble protein loss, disrupts epitopes and damages tissue integrity^55^. We thus further optimized the tissue permeabilization condition using Triton X-100 for additional functional studies, which yielded slightly improved detection efficiency (Extended Data Fig. 8a, b), and enabled simultaneous imaging of RNAs and proteins *in situ* (Fig. 6a). However, for most applications in cell lines and tissues (Fig. 1-5), methanol permeabilization was recommended, as it’s more suitable for long-term sample storage.

**Fig. 6.**
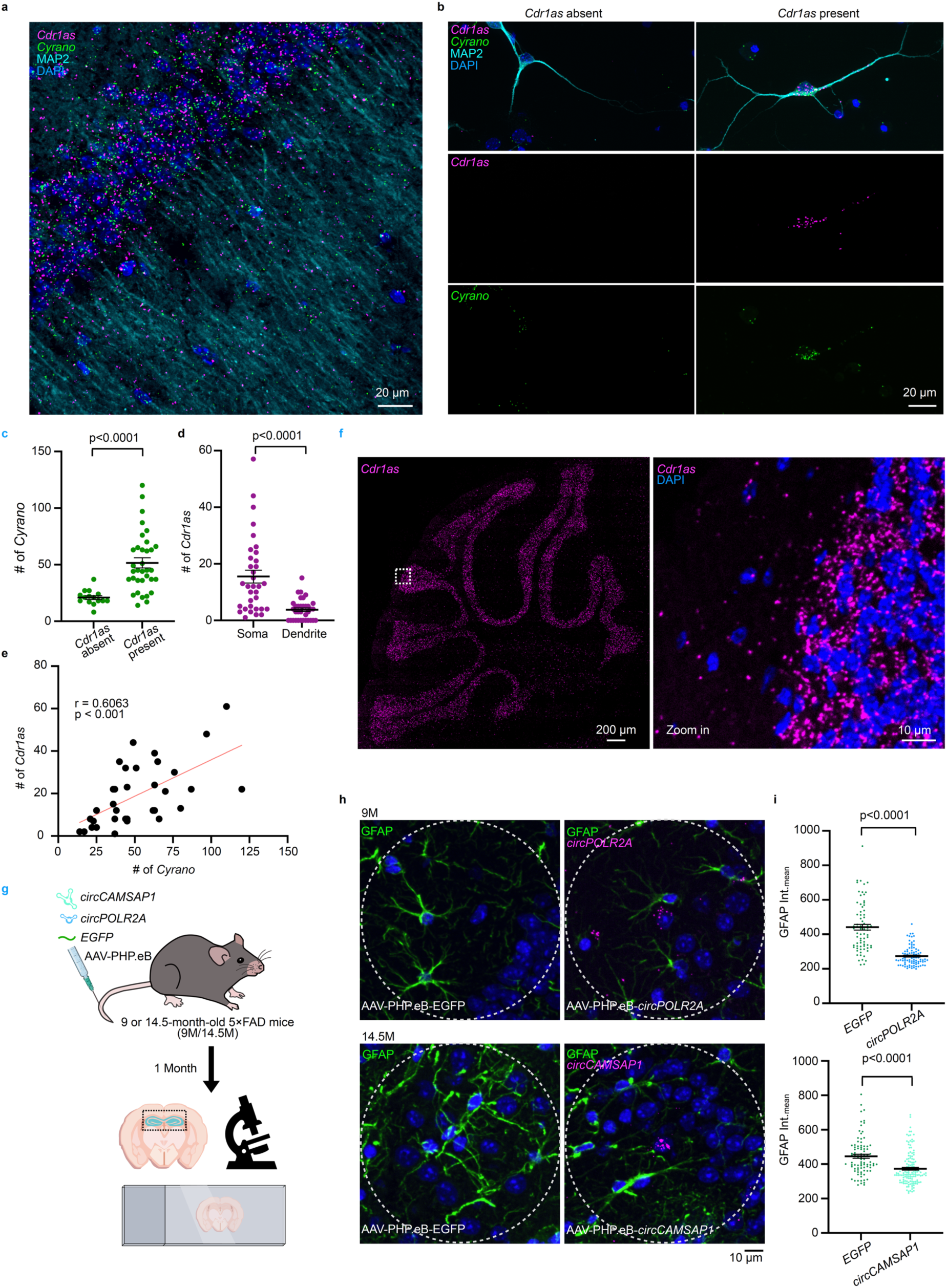
Single-molecule circRNA imaging for functional studies. **a,** Representative images of simultaneous multiplexed abcFISH imaging of *Cyrano* and *Cdr1as* together with MAP2 immunostaining in mouse hippocampal CA1 region. MAP2 staining marks dendrites. Magenta, *Cdr1as* signals. Cyan, MAP2 IF signals. **b,** Representative images of detecting *Cyrano* and *Cdr1as* in primary hippocampal neurons. *Cdr1as* was undetectable in neurons expressing low levels of *Cyrano* (left panels), but observed in neurons with high *Cyrano* expression (right panels). Green, *Cyrano* signals. Magenta, *Cdr1as* signals. Cyan, MAP2 IF signals. **c,** Statistics of *Cyrano* copy number per cell in *Cdr1as* absent or present cells shown in **b**. Data represent mean ± s.e.m.; n = 15 (*Cdr1as* absent) and 34 (*Cdr1as* present) cells; *P* values were calculated using two-tailed Student’s t-test. **d,** Statistics of subcellular distribution of *Cdr1as* in soma versus dendrites. Data represent mean ± s.e.m.; n = 34 cells; *P* values were calculated using two-tailed Student’s t-test. **e,** Positive correlation between *Cdr1as* and *Cyrano* copy numbers at single-neuron resolution. Each point represents one cell; n = 34 cells; Pearson’s r and *P* value are indicated. **f,** Representative images of *Cdr1as* detected by abcFISH in FFPE sagittal brain sections from adult mouse. *Cdr1as* shows abundant localization in the cerebellum. **g,** Schematic overview of intravenous injection and sample collection of 5×FAD mice treated with therapeutic ivcRNAs. AAV-PHP.eB vectors expressing engineered ds-cRNAs or *EGFP* control were delivered via intravenous injection. Brains were collected one month post-injection for analysis by abcFISH and immunofluorescence. **h,** Representative images of simultaneous detection of ds-cRNAs and GFAP in mouse brain sections from injected mouse. The astrocytes were labeled by GFAP IF. White dashed circles indicate examples of the 80-pixel radius regions used for GFAP mean intensity quantification. Green, GFAP IF signals. Magenta, ds-cRNA signals. **i,** Statistics of GFAP immunofluorescence intensity within an 80-pixel radius surrounding ds-cRNA-positive cells shown in **h**. For the GFP control samples, where the therapeutic circular RNAs were not present, measurement areas were selected unbiasedly. The investigator was blinded to the group allocation during data analysis. The GFAP intensity significantly decreased with ds-cRNAs delivered compared to the *EGFP* control group. Data represent mean ± s.e.m.; For 9-month-old (9M) mice, n = 74 (*EGFP*) and 91 (*circPOLR2A*) regions; for 14.5-month-old (14.5M) mice, n = 91 (*EGFP*) and 138 (*circCAMSAP1*) regions; *P* values were calculated using two-tailed Student’s t-test.

In the hippocampal CA1 region, both *Cdr1as* and *Cyrano* predominantly localized in neuronal somata, with sparse signals detected in dendrites (Fig. 6a). Single-cell quantification in isolated hippocampal neurons revealed that *Cdr1as* was not uniformly expressed across all neurons but was rather significantly enriched in cells with high *Cyrano* expression (Fig. 6b, c). Both RNAs were primarily localized in the soma in hippocampal primary neurons (Fig. 6d), and their expression levels showed significant positive correlation (Fig. 6e), supporting the model that *Cyrano* stabilizes *Cdr1as* through miR-7 destabilization^31^.

### abcFISH is compatible with FFPE section samples

The altered expression of circRNAs has been found in diseases, and their highly stable nature makes them potential biomarkers^56^. Thus, we evaluated whether abcFISH is compatible with FFPE sections, a common clinical sample format. Using abcFISH, we observed the expected cerebellum-enriched localization of *Cdr1as* (Fig. 6f), consistent with previous reports shown in canonical *in situ* hybridization^11^, supporting the method’s potential for future translational and diagnostic applications.

### Therapeutic ds-cRNAs reduce astrocyte infiltration in the hippocampus of AD animals

Recently, therapeutic ds-cRNAs have shown efficacy in dermatological and neurological disease models^30, 32^. Conventional assays lack the ability to correlate circRNA presence and local protein response at single-cell resolution. abcFISH overcomes this limitation by enabling simultaneous detection of circRNAs and proteins in intact tissues (Fig. 6a).

We delivered AAV-PHP.eB vectors expressing ds-cRNAs: *circPOLR2A(9,10)*, *circCAMSAP1(2,3)*, or the *EGFP* control via intravenous injection to the 5×FAD mice and analyzed brain sections one month later (Fig. 6g). Using abcFISH, we detected exogenous ds-cRNAs in the hippocampus and performed concurrent immunofluorescence (IF) for astrocytes (Fig. 6h). Quantification of GFAP intensity within an 80-pixel radius around ds-cRNA positive cells (Fig. 6h) exhibited a 38% and 16.4% reduction in astrocyte infiltration around *circPOLR2A(9,10)-* and *circCAMSAP1(2,3)-*expressing cells, respectively, compared to controls (Fig. 6i). These findings directly showed the efficacy of ds-cRNA aptamers in reducing neuroinflammation mediated by astrocyte infiltration, confirming their local anti-inflammatory effects *in vivo*.

## Discussion

CircRNAs have traditionally been studied using bulk RNA sequencing, an approach without their spatial context and cellular heterogeneity. Advances in imaging have enabled high-resolution visualization of mRNAs, lncRNAs and modified RNAs^25, 26, 36, 57–59^, however, visualizing circRNAs remains challenging due to their low abundance and sequence overlap with linear isoforms^16, 28^. Here we developed abcFISH (Fig. 1a-e) as a robust method for multiplexed, single-molecule imaging of circRNAs in fixed cells and tissues (Fig. 2 and 3). By combining BSJ-specific probing with RCA, abcFISH outperformed existing methods with its high SNR, high specificity, high sensitivity and low cost (Fig. 1f-h and Extended Data Fig. 3a, 4). A major advantage of abcFISH is its ability for the direct quantification of circRNA abundance and subcellular distribution, eliminating interference from linear isoforms. This approach simultaneously preserves native spatial context and decreases experimental costs.

Leveraging this capability, we resolved several questions regarding circRNA localization and regulation. We visualized multiple circRNAs and circRNA alternative circularization simultaneously, providing the direct evidence to show their distinct cellular localization (Fig. 2). Furthermore, we imaged the heterogenous expression of *circPOLR2A(9,10)* across tissues and hippocampal cell types in a KI mouse model (Fig. 3, 4). The high expression of this human circRNA cassette observed in mouse tissues provides direct evidence supporting the requirement of strong intronic pairing for circRNA biogenesis^17^. By integrating spatial profiles with RBPs known to control circRNA turnover^46^, our analyses indicated a potential combinatorial RBP effect on modulating circRNA abundance in the nervous system (Fig. 4). Interestingly, both abundant *circAnks1b(5,6,7,8)* and *Cdr1as* exhibited distinct subregional and subcellular localization patterns in the brain (Fig. 5 and Extended Data Fig. 7), indicating their potential roles await for functional studies. For ncRNAs with known functions, we calculated the positive correlation between *Cyrano* and *Cdr1as* levels at the single-cell level, providing the direct visual evidence supporting the model of *Cyrano*-mediated stabilization of *Cdr1as*^31^ (Fig. 6a-e). abcFISH was compatible with freshly prepared FFPE tissues (Fig. 6f) and enabled measurement of the efficacy of therapeutic ds-cRNAs *in vivo* by directly visualizing the delivered ds-cRNAs and the reduction of astrocyte infiltration surrounding ds-cRNA-positive regions (Fig. 6g-i), collectively showing its potential for future circRNA-based therapeutics and diagnostics. Altogether, applications of abcFISH in these examined contexts in cells and *in vivo* (Fig. 2-6) have established it as a robust and user-friendly method for circRNA spatial detection. Given the increasing evidence that circRNAs localize to various subcellular compartments, including the nucleus^60^, mitochondria^61^ and P-bodies^62^, abcFISH is also positioned for functional studies in details. Moreover, integration with proximity labeling approaches^51^ could complement abcFISH to dissect local interactions of circRNAs within specific organelles. Future integration of abcFISH with other approaches for circRNA studies will provide additional insights into functional understanding of circRNAs, mirroring advances in mRNA and lncRNA research^63, 64^.

Despite these advantages, abcFISH has some limitations. First, as with all FISH-based techniques, its detection efficiency depends on RNA accessibility, requiring optimization of permeabilization and protease digestion. We recommend adjusting the probing region flanking the BSJ to enhance hybridization. Second, while sufficient for multi-target experiments, the multiplexing capacity is limited compared to current mRNA *in situ* sequencing methods^36, 65^, given that circRNAs are less abundant and harbor only one amplicon per BSJ site. Further efforts are warranted to expand multiplexity by employing additional fluorophores or sequential imaging with protocol compatible with low abundant circRNAs. Third, although abcFISH was compatible with freshly prepared FFPE tissues (Fig. 6f), its performance on archival FFPE specimens with different storage durations has not been evaluated. Fourth, although several RBPs were likely to regulate circRNA turnover in mouse brain, their regulatory roles need to be verified. Finally, while abcFISH provides high-resolution spatial information, it does not capture the circRNA dynamics in live cells, and the displacement of the RCA amplicons from the original circRNA localization should be considered during colocalization analysis.

## Acknowledgments

We acknowledge X.-L. Xu from CEMCS, CAS for support on microscopy; and G.-H. Yuan, S.-N. Zhai, X. Feng and M.-Y. Wei for supporting our experiments. This work was supported by the National Key R&D Program of China (2021YFA1300501), the Strategic Priority Research Program of the Chinese Academy of Sciences (XDB0570000), Science and Technology Commission of Shanghai Municipality (23DX1900100; 23DX1900101) to L.-L.C.

This work has been supported by the New Cornerstone Science Foundation through the New Cornerstone Investigator Program to L.-L.C and supported by the Ruisi Research Center for Life Science, Minhang District, Shanghai. L.-L.C. is also a SANS Senior Investigator.

## Author contributions

L.-L. C. conceived the project. Y.-X. L. and B.-W. J. designed and performed experiments with the help of K.-M. L.; Y.-X. L., B.-W. J., K.-M. L. and L.-L. C. wrote the manuscript. L.-L. C. supervised the project.

## Methods

### Cell culture and sample preparation

Human HeLa cells were purchased from the American Type Culture Collection (ATCC; http://www.atcc.org). HT29 and DLD-1 cells were purchased from National Collection of Authenticated Cell Cultures. HeLa cells were maintained in DMEM supplemented with 10% Fetal Bovine Serum (FBS). HT29 and DLD-1 cells were maintained in RPMI-1640 supplemented with 10% FBS. All cell lines were cultured at 37 °C with 5% CO₂, and were routinely screened to exclude mycoplasma contamination.

Poly(I:C) (1 μg/mL, Sigma, P9582) transfection was carried out using Lipofectamine 2000 Reagent (Thermo Fisher Scientific–Invitrogen, 11668019) into cells for 4 hours, then cells were treated with 4% paraformaldehyde (PFA) for fixation.

To prepare cell samples, cells cultured in 18-well chambered cover glass (Cellvis, C18SB-1.5H) with 60%-80% confluency were washed with DPBS twice and fixed with 4% PFA for 10 min at room temperature. After fixation, cells were washed three times with DPBS. Then, the cells were permeabilized with either Triton X-100 in smFISH or cold methanol at −20 °C for 1 hour in abcFISH and ampFISH.

### abcFISH

Fixed samples permeabilized by either Triton X-100 or methanol were equilibrated by equilibration buffer (0.1 mg/mL yeast tRNA (Solarbio, T8630), 0.4 U/µL RNasin (Vazyme, R301-03), 100 mM glycine, 0.1% Tween-20 and 2 mM ribonucleoside-vanadyl complexes (Beyotime, R0107) in RNase-free 1×PBS (Thermo Fisher Scientific–Invitrogen, AM9624)). Phosphorylated padlock probes (100 nM per probe) and primer probes (100 nM per probe) in hybridization buffer (10% formamide, 2× SSC, 0.1 mg/mL yeast tRNA, 0.2 U/µL RNasin, 20 mM ribonucleoside-vanadyl complexes) were added to cell samples or pretreated brain slices and incubated overnight at 40 °C. After hybridization, samples were washed twice with PBSTR (0.1% Tween-20, 0.2 U/µL RNasin in RNase-free 1×PBS) at 37 °C for 20 min each time, followed by washing with 4× SSC buffer in PBSTR once at 37 °C for 20 min. After another quick wash with PBSTR, the ligase reaction mixture (0.25 U/μL T4 DNA ligase (Thermo Scientific, EL0012), 0.5 mg/ml BSA (Sangon, A000290), and 0.4 U/μL of RNasin in 1× T4 DNA ligase buffer) was then added to the sample for a 2 h incubation at 25 °C. Following ligation, samples were washed with PBSTR twice and incubated with RCA reaction solution (0.5 U/μL Phi29 DNA polymerase (Vazyme, N106-02), 250 μM dNTP (TaKaRa, 4019), 0.5 mg/mL BSA and 0.4 U/μL of RNasin in 1× Phi29 buffer) at 30 °C for 2 h. After double washing with PBSTR, the samples were detected by the fluorescent probes (100 nM per probe in hybridization buffer) for 1 hour at room temperature. The excess fluorescent probes were removed with hybridization buffer three times at room temperature, 10 min each. The cells were incubated with 1 µg/mL DAPI (Thermo Fisher Scientific–Invitrogen, D1306) diluted in 2× SSC for 1 min at room temperature for nuclear staining. For staining tissue structures, 100 nM TRITC Phalloidin (Yeasen, 40734ES75) was added during detection. The samples were then mounted in Slowfade Diamond antifade mounting medium (Thermo Fisher Scientific–Invitrogen, S36963).

For combination of immunofluorescence and abcFISH in Fig. 6 and 7, fixed samples were permeabilized with 0.5% Triton X-100 for 10 min, and were washed with PBSTR three times for 5 min each time. Then the samples were blocked with 1% BSA for 1 h at room temperature. Primary antibodies were diluted with 1% BSA (MAP2, Epizyme, R013804, 1:200 ; GFAP, CST, 3670S, 1:500) and incubated for 1 h at room temperature. After washing three times with PBSTR, fluorescent secondary antibodies (Goat anti-Mouse Secondary Antibody Alexa Fluor 488 or Goat anti-Rabbit Secondary Antibody Alexa Fluor 555, Invitrogen, A-11001 and A-21428) were diluted 1: 1,000 in 1% BSA and incubated for 1 h at room temperature. After washing three times with PBSTR, the samples were equilibrated and hybridized with probes following abcFISH procedures.

For abcFISH probe design, a 56 nt “targetable zone” centered on the BSJ is proposed (Extended Data Fig. 1h). All abcFISH probes in this study (except those used in Fig. 1b) were centered at position “0”. Probe sequences are listed in Supplementary Table 1.

For fluorophore selection, AF488, AF568 and AF647 all perform well in cells, whereas AF647 is optimal for tissues due to reduced autofluorescence in the far-red channel.

### smFISH

All smFISH probes were designed using Stellaris Probe Designer and labeled with either Gold550 or Red650 at the 3′ end. smFISH was carried out as described previously^66^. In brief, the PFA-fixed cells were permeabilized with 0.5% Triton X-100 for 10 min. Cells were incubated in 10% formamide/2× SSC for 10 min at room temperature followed by hybridization at 37 °C for 16 h. After hybridization, the cells were washed twice with 10% formamide/2× SSC at 37 °C, each time for 30 min. The cells were incubated with 1 µg/mL DAPI diluted in 2× SSC for 1 min at room temperature for nuclear staining. The samples were then mounted in SlowFade Diamond antifade mounting medium. A list of the probe sequences for smFISH is provided in Supplementary Table 1.

### ampFISH

ampFISH was carried out as described previously^28^. In brief, PFA-fixed cells were permeabilized by subsequently cold methanol at -20 °C for 1 hour and equilibrated with 10% formamide/2× SSC for 10 min at room temperature. Then, the hybridization reaction initiated at 37 °C overnight with 50 µL hybridization mixture: 10% dextran sulfate (Sigma-Aldrich, s4030), 1 mg/mL tRNA, 2 mM RVC, 0.02% BSA, 10% formamide, 2× SSC and 2 µL each of donor and acceptor probes (10 ng/µL). After hybridization, the cells were washed twice with 10% formamide/2× SSC at room temperature, each time for 10 min. Subsequently, HCR was performed in HCR buffer that consisted of 5× SSC, 10% dextran sulfate, 0.05% Tween-20, and 125 nM of each HCR hairpin at 25 °C for 4 h. After HCR reaction, the cells were washed twice with 10% formamide/2× SSC at room temperature, each time for 10 min. The cells were incubated with 1 µg/mL DAPI diluted in 2× SSC for 1 min at room temperature for nuclear staining. The samples were then mounted in SlowFade Diamond antifade mounting medium. A list of the donor probes, acceptor probes and HCR hairpins is provided in Supplementary Table 1.

### SplintR-RCA

SplintR-RCA was carried out as described previously^28^. In brief, 100 nM padlock probes were hybridized in 6× SSC and 10% formamide for 2h at 37 °C. After washing with PBSTR three times, SplintR ligation mix (0.5 U/mL SplintR ligase (NEB, M0375S), 0.2 mg/ml BSA (Sangon, A000290), and 0.4 U/μL of RNasin in 1× SplintR ligase buffer) was added for ligation at 37 °C for 1 h, followed by three rinses with PBSTR. Prior to RCA, 100 nM primer probes were added in 2× SSC and 20% formamide at 37 °C for 30 min. After three rinses with PBSTR, RCA reaction solution mentioned above was added at 30 °C for 2 h. After double washing with PBSTR, the samples were detected by the fluorescent probes for 1 hour at room temperature. The excess fluorescent probes were removed with hybridization buffer three times at room temperature. The cells were incubated with DAPI solution for nuclear staining and were finally mounted in Slowfade Diamond antifade mounting medium. Probe sequences for SplintR-RCA are provided in Supplementary Table 1.

### BaseScope assay

A BaseScope probe targeting *circHIPK3(2*) was designed by Advanced Cell Diagnostics (715871). BaseScope assays were performed using a BaseScope™ Reagent Kit v2 RED (Advanced Cell Diagnostics, 323900) according to the manufacturer’s instructions.

### Knockdown with siRNAs

For *circHIPK3(2)* knockdown with siRNAs, HeLa cells were seeded in 24-well plates, then siRNA (GenScript) was transfected at a final concentration of 10 nM using RNAiMax (Thermo Fisher Scientific–Invitrogen, 13778030) 48 h before the passage.

### RNA isolation and RT-qPCR

Total RNAs were extracted with Trizol Reagent (Ambion, 15596018). The reverse transcription and RT-qPCR were carried out using PrimeScript RT Master Mix (TaKaRa, RR036A) and SYBR Green Real-time PCR Master Mix (TOYOBO, QPS-201C). All primers used for RT-qPCR were listed in Supplementary Table 1.

### Plasmid construction

To generate standard curves for the absolute quantification of circRNAs via qPCR, plasmid constructs containing the exact back-splicing junction (BSJ) sequence of each target circRNA were created. For each circRNA (including *circASPH(2,3)*, *circCDYL(2)*, *circSPECC1(4)*, *circHIPK3(2)* and *circPOLR2A(9,10)*), a DNA fragment spanning the entire circularized exon(s) with the BSJ positioned centrally was amplified from cDNA using PrimerSTAR DNA polymerase. The purified PCR products were subsequently cloned into pUC57 plasmids (Transgen, CU201-03). Sequencing-validated plasmids were extracted using Mini-Prep kit (TIANGEN, DP103). The primers used for plasmid construction were listed in Supplementary Table 1.

### qPCR measurement of circRNA copy number

A serial dilution of purified plasmid harboring circRNA-forming exons was used for qPCR to generate a standard curve. The copy number of the diluted DNA template was calculated by DNA/RNA Copy Number Calculator from the following website (http://endmemo.com/bio/dnacopynum.php). To measure the circRNA copy per cell, total RNAs extracted from 1 × 10^6^ HeLa cells were reversely transcribed into cDNAs, and aliquots of cDNAs from either 5000 HeLa cells were further used for qPCR. The copy numbers of circRNAs were quantified from the standard curve.

### Deparaffinization of FFPE sections

Two-month-old WT mice were euthanized, and the brains were collected and fixed in neutral buffered formalin for at least 48 h, followed by paraffin embedding. Brain sagittal sections of 2 μm thickness were prepared for subsequent analyses.

Sections were deparaffinized in xylene twice (5 min each at room temperature), then incubated in ethanol twice (2 min each). Sections were then subjected to antigen retrieval by immersing in 1× citrate buffer (Beyotime, P0081) at 95°C for 15 min, followed by cooling to room temperature. After two washes in distilled water, sections were storaged in cold methanol at -80°C until use.

### Experimental animals

The following mice were used in this study: C57BL/6, 5×FAD and C_OE (circRNA-overexpression) mice. The 5×FAD mice on a C57BL/6J background were provided as gifts by N. Jing at the Guangzhou Institute of Biomedicine and Health, Chinese Academy of Sciences (CAS). Transgenic mice with constitutive ds-cRNA overexpression were established on a C57BL/6 background. In brief, a construct was inserted into the ROSA26 locus containing EF1α promoter, the exonic sequences destined for circularization from human *circPOLR2A(9,10)*, and flanking intronic regions harboring reverse-oriented complementary sequences. Correct insertion was verified by Southern blot analysis combined with restriction enzyme digestion. Embryonic stem cell clones with accurate targeting were injected into C57BL/6 blastocysts to produce C_OE mice. Genotyping was performed by PCR amplification of genomic DNA fragments from tail biopsies using specific primer sets.

### Tissue collection and sectioning

The brain tissue was collected after rapid decapitation. Fresh brain tissues were immediately snap-frozen in liquid nitrogen and stored at −80 °C until sectioning. For cryosectioning, frozen brains were mounted onto the specimen holder with OCT (Richard-Allan Scientific, 6502), and coronal sections were cut at a thickness of 10 µm using a cryostat (RWD, Minux FS800) maintained at −20 °C. Sections were mounted onto Superfrost™ Plus slides (Epredia, 4951PLUS-001E) and stored at −80 °C until further use.

To prepare tissue samples for abcFISH, tissue sections were washed with DPBS once and fixed with 4% paraformaldehyde for 15 min at room temperature. After fixation, tissues were washed three times with DPBS. Then, the sections were permeabilized with cold methanol for 1 hour at −80 °C for abcFISH.

### Primary hippocampal neuron culture from P0 mice

Hippocampal neurons were isolated from postnatal day 0 (P0) C57BL/6 mice of either sex. Neonates were decapitated, and the brains were rapidly removed and transferred to ice-cold Hank’s Balanced Salt Solution (HBSS; without Ca²⁺/Mg²⁺, Thermo Fisher Scientific–Invitrogen, Cat# 14175079). Hippocampi were dissected under a stereomicroscope, and the meninges were carefully removed. The tissue was enzymatically digested in 10 U/mL papain (Worthington Biochemical Corporation, LS003126) and 0.1 mg/mL DNase I (Sigma, DN25) for 18 min at 37 °C with gentle agitation, followed by washing in HBSS. Digestion was terminated by adding an equal volume of Dulbecco’s Modified Eagle Medium (DMEM; Gibco) supplemented with 10% fetal bovine serum (FBS). The tissue was then mechanically dissociated by gentle trituration using a 1 mL pipette tip, followed by centrifugation at 1000 rpm for 5 min.

The resulting cell pellet was resuspended and plated in DMEM/F12 medium supplemented with 10% FBS, 1 mM sodium pyruvate (Thermo Fisher Scientific–Invitrogen, 11360070), 2 mM L-glutamine (Thermo Fisher Scientific–Invitrogen, 25030081), and 1% penicillin/streptomycin, onto 35-mm glass-bottom dishes coated with poly-D-lysine (0.01 mg/mL; Sigma). After 12 h, the culture medium was replaced with Neurobasal medium (Gibco) supplemented with 2% B27 (Gibco) and 2 mM L-glutamine. Neurons were fixed for staining at 2–3 days in vitro (DIV). Cultures were maintained at 37 °C in a humidified 5% CO₂ incubator.

### Animal use and care

Mice were bred and maintained under specific-pathogen-free (SPF) conditions at the Center for Excellence in Molecular Cell Science (CEMCS), CAS. All animal experiments were approved by the Committee of Use of Laboratory in the Center for Excellence in Molecular Cell Science, Chinese Academy of Sciences. Mice were raised together with littermates in pathogen-free environment, and their health status was routinely checked. No more than 5 mice were housed in one cage. The mice were housed in standard cages with free access to food and water. Mice were maintained in a 12-h light/dark cycle at 22–26 °C.

### Statistics analysis

The data used in this study are presented as mean ± s.d. or s.e.m. Statistical analyses (two-tailed Student’s t-test, one-way analysis of variance (ANOVA), linear regression, correlation analysis and so on) were performed using existing software (GraphPad Prism9). P < 0.05 was considered as significant.

### Image acquisition

Confocal microscope images were acquired using the Nikon AX/AX R microscope equipped with NSPARC and Nova-SD spinning disk, and operated with NIS-Element Software.

For cultured cells, images were captured at a z-interval of 0.3 µm per slice and at 60× magnification. For tissue sections, images were captured at a z-interval of 0.5 µm per slice and at 40× magnification.

### Image analysis and quantifications

Spot detection, cell segmentation and SNR analysis were performed with Big-FISH (https://github.com/fish-quant/big-fish), a Python implementation of FISH-Quant^67^. It starts with a Laplacian of Gaussian filter (LoG) to enhance the spot signal. A local maximal algorithm localizes every peak in the image, then a threshold is applied to discriminate the actual spots from the nonspecific background signal. The nuclei were segmented with a pretrained Unet model and the cells were segmented with a watershed algorithm. Detected spots were subsequently assigned to segmentation results. For brightfield images generated from BaseScope, circRNA signals were detected with Icy(v 2.5.4.0)^68^.

The SNR was calculated using the following formula: SNR = P_signal_ / P_background_ = (maximum intensity of spots signal – mean intensity of background) / s.d. of background signal, in which P_signal_ means the difference of maximum signal intensity and background mean intensity, and P_signal_ means the standard deviation of background intensity. Background is a region twice larger surrounding the spot region.

### Analyses of circRNA expression and circularization efficiencies

FPB_circ_ of circRNAs detected in HeLa cells and FPB_linear_ of the corresponding linear RNAs were obtained from previous sequencing data^6^. The circularization efficiencies for *circPOLR2A(9,10)* in H9 cells and human kidney samples were calculated as FPB_circ_ / (FPB_linear_+ FPB_circ_), using data retrieved from CIRCpedia V3^69^.

### eCLIP-seq analyses

eCLIP-seq datasets were obtained from the ENCODE database (https://www.encodeproject.org/)^70^ and analyzed using version 2.2 of previously published eCLIP-seq processing pipeline^71^. Specifically, RNA-seq reads were first trimmed by Cutadapt (v1.14; parameters: --match-read-wildcards --times 1 -e 0.1 -O 5 --quality-cutoff 6 -m 18) and then mapped to ribosomal DNA sequences using Bowtie2 (v2.2.5; parameters: default) to remove reads mapped to rRNAs. Next, the remaining high-quality reads were aligned to the hg38 reference genome using STAR (v2.5.2b; parameters: --outFilterMultimapNmax 30 --outFilterType BySJout --outFilterMultimapScoreRange 1 --outFilterScoreMin 10 --outFilterMismatchNoverLmax 0.01). Unmapped reads were subsequently aligned to a custom reference consisting of concatenated duplicated *circXPO1(2,3,4)* sequences. PCR duplicates were then removed using barcodecollapsepe.py from Yeo lab eCLIP-seq processing pipeline^71^. Reads spanning the BSJ site were subsequently extracted by SAMtools (v1.6). These BSJ-spanning reads were then utilized for the identification of potential RBP binding peaks by Clipper3^72^.

### Screening of candidate RBPs associated with circRNA expression heterogeneity

Gene expression matrix and cell cluster annotation of mouse hippocampus in 10×v3 Single-cell RNA-seq datasets were downloaded from Allen Brain Atlas^50^. Correlation analysis between RBP expression levels and circRNA imaging-derived abundances across distinct hippocampal cell types was performed to identify candidate RBPs potentially associated with circRNA expression heterogeneity in the mouse hippocampus from circRNA previous research. Specifically, Spearman’s rank correlation was calculated for each RBP previously implicated in circRNA biogenesis^46^ by Python (v3.13.0) library SciPy (v1.15.2). Subsequently, false discovery rate (FDR) correction was applied to the resulting p-values to account for multiple testing by Python (v3.13.0) library statsmodels (v0.14.4). RBPs showing statistically significant correlations (FDR adjusted p < 0.05) were considered as candidate regulators associated with circRNA expression heterogeneity.

## Data availability

All data supporting the findings of this study are available in the Article. Source data are provided with this paper.

## Supplemental material

### Supplemental Table 1

Sequence details used in this study, four sections are included: abcFISH probe sequences, ampFISH probe sequences, SplintR-RCA probe sequences, smFISH probe sequences, siRNA sequences and primer sequences.

## Extended Data Figures and Figure legends

**Extended Data Fig. 1.**
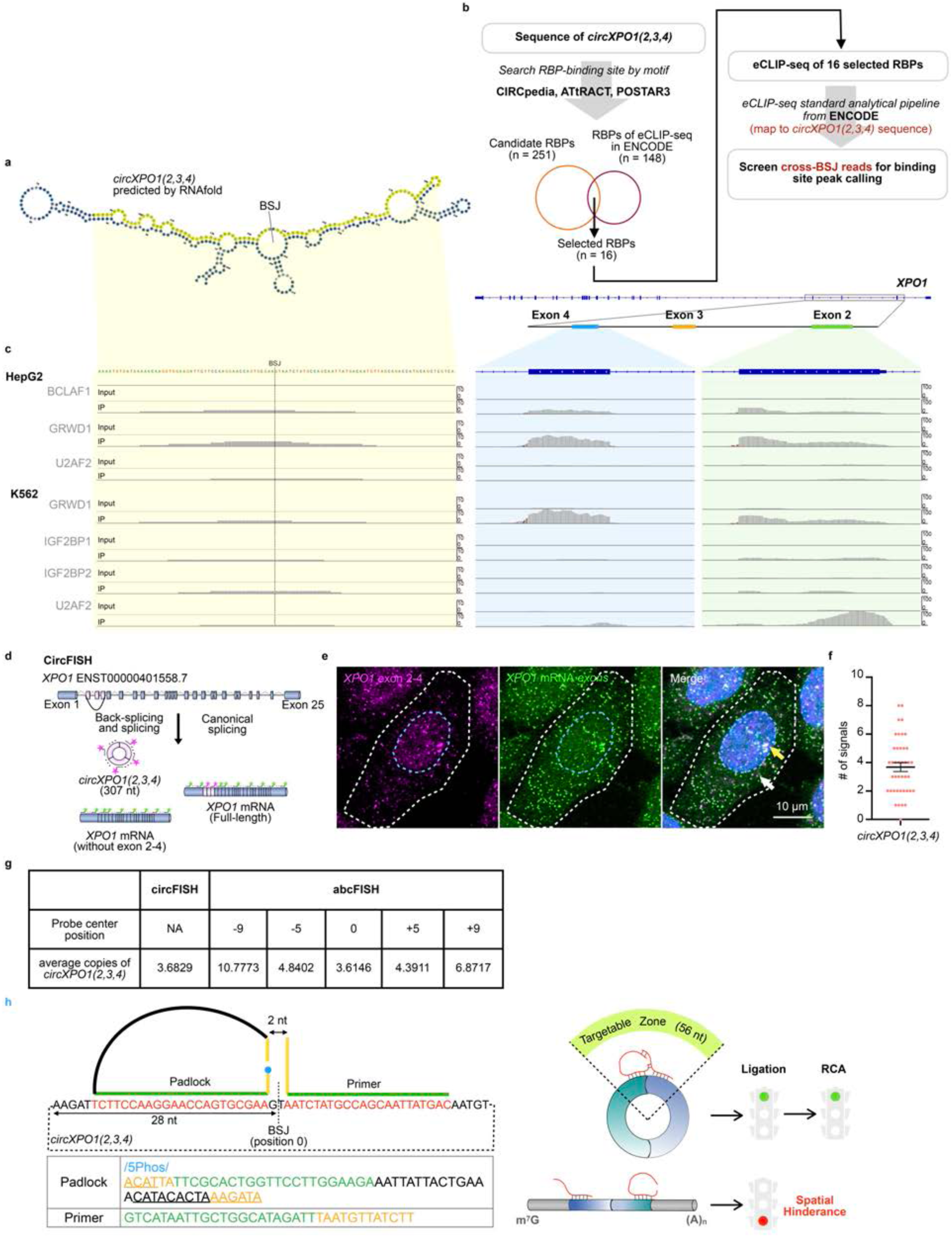
Predicted structure, RBP binding and smFISH verification of circXPO1(2-4) **a,** Predicted RNA secondary structures of *circXPO1(2,3,4)* generated by RNAfold, showing stem-loops flanking the BSJ site (marked by a black line). The BSJ site is indicated by the black line. Sequences 50 nt upstream and downstream of BSJ are highlighted in yellow. **b,** Workflow for analyzing published eCLIP-seq data to identify RBPs associated with *circXPO1(2,3,4)*. circRNA sequence and RBP binding information were obtained from CIRCpedia, ATtRACT, and POSTAR3 databases. Sixteen RBPs overlapping with ENCODE eCLIP-seq datasets were selected, and cross-BSJ reads were mapped to a 100-nt region centered on the BSJ. **c,** RBPs mapped to the *circXPO1(2,3,4)* BSJ region (the left panel) and linear *XPO1* exon 2 and 4 (right panels), based on eCLIP-seq data from HepG2 and K562 cell lines, showing less RBP binding on circRNA than linear RNA. **d,** Schematic of the circFISH probe design for distinguishing *circXPO1(2,3,4)* from linear *XPO1* transcripts. Gold550-labeled probes target exons 2–4 (shared by both linear and circular forms), while Red650-labeled probes hybridize to exons found only in linear *XPO1* mRNA. **e,** Representative circFISH images of *circXPO1(2,3,4)* in HeLa cells. The transcription site is indicated by a yellow arrow, and a circRNA-specific focus is marked by a white arrow. The nuclear boundary is outlined with a blue dashed line; the cellular boundary is indicated with a white dashed line. Magenta, exons 2–4 signals (Gold550); Green, linear-specific *XPO1* exon signals (Red650). **f,** Statistics of Gold550-only puncta per cell from circFISH shown in **e**. Data represent mean ± s.e.m.; n = 41 cells. **g,** Comparison of *circXPO1(2,3,4)* signal numbers between circFISH and abcFISH. abcFISH probes spanning the BSJ (−5 to +5 nt) showed comparable performance to circFISH, confirming high specificity and quantitative accuracy. **h,** Schematic of abcFISH probe structures using *circXPO1(2,3,4)* as an example (left). The sequences flanking the BSJ are indicated. Sequences targeted by abcFISH probes are shown in red, and the hybridization sequences of abcFISH are shown in green. Complementary sequences between the padlock probe and the primer are highlighted in yellow. The 5′ phosphorylation modification is colored blue. Sequences recognized by the fluorescent detection probe are underlined. The right panel illustrates the targetable zone of 56 nucleotides spanning the BSJ. Due to spatial hindrance, ligation does not proceed on linear RNAs, ensuring circRNA-specific detection.

**Extended Data Fig. 2.**
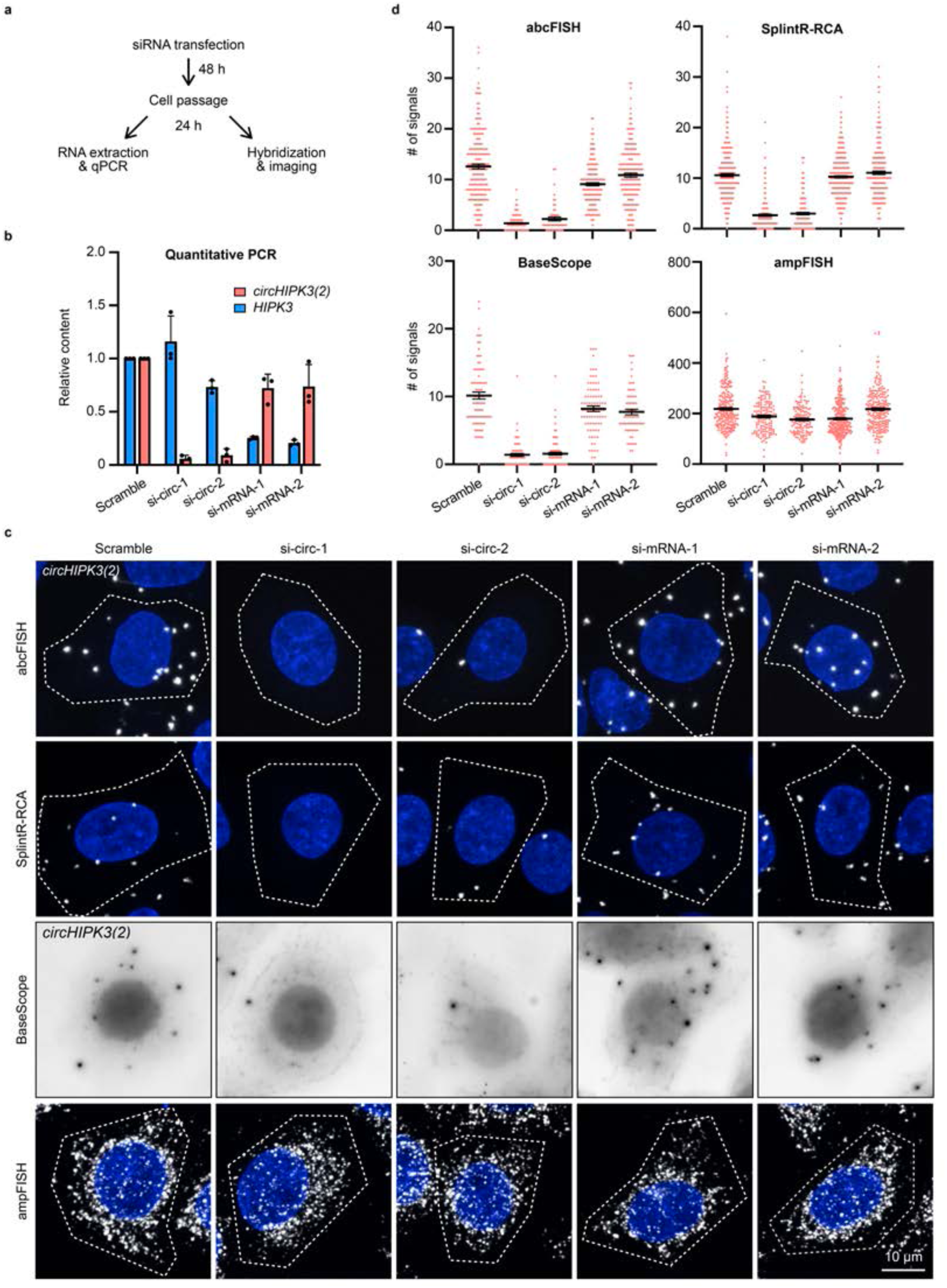
Specificity validation of direct circRNA imaging methods by siRNA knockdown. **a,** Workflow of the siRNA knockdown experiment targeting *circHIPK3(2)*. Cells were transfected with siRNAs, passaged 48 h post-transfection, then processed either for RNA extraction and qPCR or for imaging using abcFISH, SplintR-RCA, BaseScope, or ampFISH. **b,** qPCR quantification of linear and circular HIPK3 after siRNA transfection. siRNAs targeting *circHIPK3(2)* (si-circ-1 and si-circ-2) efficiently knocked down *circHIPK3(2*) expression. Data represent mean ± SD of three biological replicates. **c,** Representative images of *circHIPK3(2)* in HeLa cells using abcFISH, SplintR-RCA, BaseScope, and ampFISH following siRNA transfection. **d,** Statistics of *circHIPK3(2)* signals per cell for each method under different siRNA treatment shown in **c**. abcFISH, SplintR-RCA, and BaseScope showed a marked reduction in detected signals upon *circHIPK3(2)* knockdown, whereas ampFISH consistently produced hundreds of signals per cell regardless of siRNA treatment. Data represent mean ± s.e.m.; for abcFISH, n = 208 (scramble), 142 (si-circ-1), 52 (si-circ-2), 187 (si-linear-1), and 189 (si-linear-2) cells; for SplintR-RCA, n = 304 (scramble), 193 (si-circ-1), 185 (si-circ-2), 337 (si-linear-1), and 258 (si-linear-2) cells; for BaseScope, n = 84 (scramble), 79 (si-circ-1), 94 (si-circ-2), 81 (si-linear-1), and 66 (si-linear-2) cells; for ampFISH, n = 234 (scramble), 143 (si-circ-1), 138 (si-circ-2), 278 (si-linear-1), and 221 (si-linear-2) cells; *P* values were calculated using a one-way analysis of variance (ANOVA).

**Extended Data Fig. 3.**
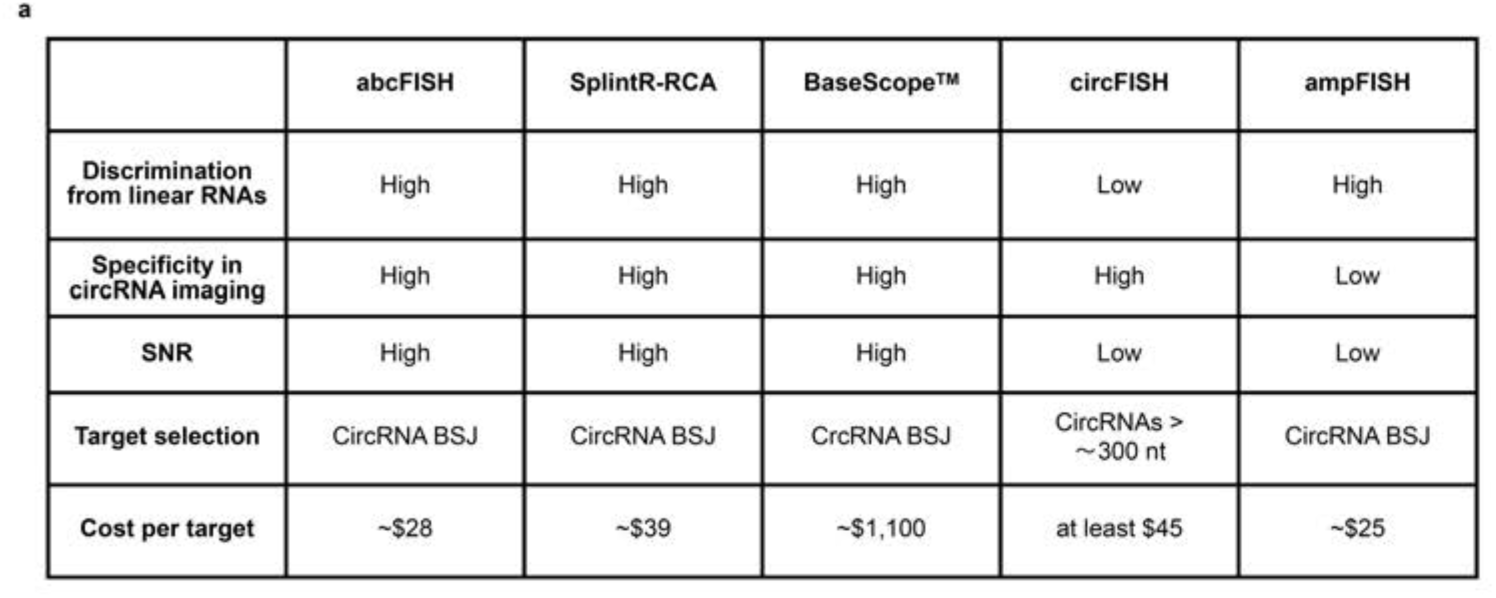
Comprehensive comparison of abcFISH with other circRNA imaging methods. **a,** Feature comparison of abcFISH, circFISH, SplintR-RCA, BaseScope, and ampFISH. abcFISH demonstrates superior overall performance, combining high feasibility, high specificity, high SNR, low cost and no circRNA size limitation.

**Extended Data Fig. 4.**
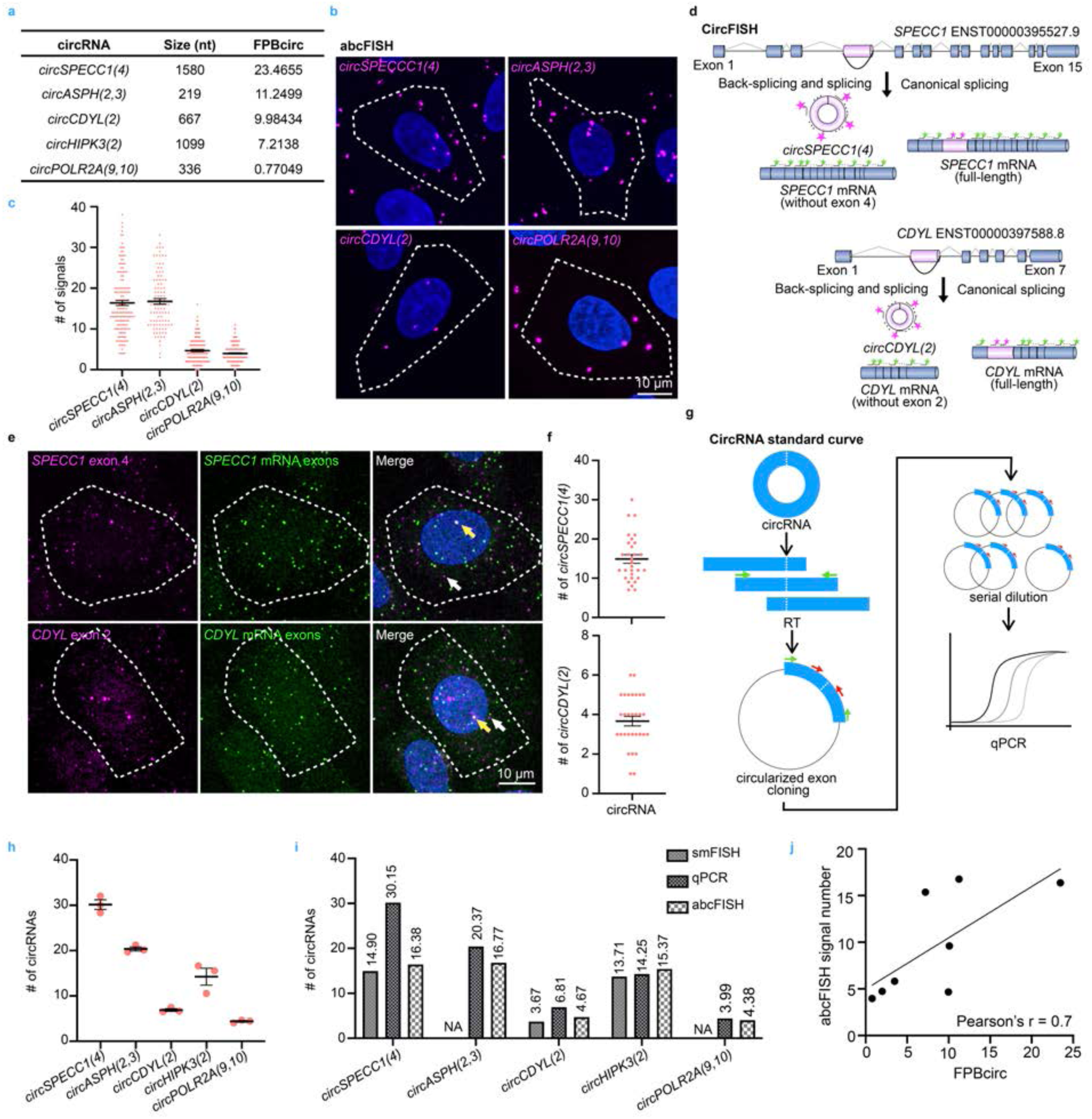
abcFISH shows concordance with established circRNA quantification methods. **a,** Overview of five circRNAs analyzed in this study, including *circSPECC1(4)*, *circASPH(2,3)*, *circCDYL(2)*, *circHIPK3(2*) and *circPOLR2A(9,10)*. Their lengths and expression levels (FPB_circ_) in HeLa cells are indicated. **b,** Representative images of abcFISH detecting *circSPECC1(4)*, *circASPH(2,3)*, *circCDYL(2)* and *circPOLR2A(9,10)* in HeLa cells. Cellular boundaries are indicated with white dashed lines. **c,** Statistics of abcFISH detected circRNA signals per cell for *circSPECC1(4)*, *circASPH(2,3)*, *circCDYL(2)* and *circPOLR2A(9,10)* in HeLa cells. Data represent mean ± s.e.m.; n = 146, 98, 130 and 181 cells, respectively. **d,** Schematic of linear and circular *SPECC1* and *CDYL* transcripts. Red650-labeled probes target exon 4 of *SPECC1* and exon 2 of *CDYL* (shared by both linear and circular forms). Gold550-labeled probes hybridize to exons found only in linear mRNA. **e,** Representative circFISH images of *circSPECC1(4)* and *circCDYL(2)* in HeLa cells. The transcription site is indicated by yellow arrows, and circRNA-specific foci are marked by white arrows. The cellular boundaries are indicated with white dashed lines. Magenta, *SPECC1* exon 2 signals or *CDYL* exon 2 signals (Red650); Green, linear-specific exon signals (Gold550). **f,** Statistics of circFISH signals per cell for *circCDYL(2)* and *circSPECC1(4)* in HeLa cells. Data represent mean ± s.e.m.; n = 30 cells per group. **g,** Workflow for generating plasmid standards for absolute qPCR quantification. cDNA containing circularized exons was reverse-transcribed, amplified with primers (green) flanking the BSJ, and cloned into pUC57. Serial dilutions of plasmids were used to generate standard curves with qPCR primers (red). **h,** Absolute quantification of circRNA copy number per HeLa cell by qPCR. Data are represented as with mean ± SD of three biological replicates. **i,** Comparison of circRNA copy number per cell measured by abcFISH, circFISH, and qPCR for *circCDYL(2)*, *circSPECC1(4)*, *circHIPK3(2)* and *circPOLR2A(9,10)*. Mean values are indicated above bars. **j,** Positive correlation between circRNA signal numbers (from panel **c** and Fig. 2i) and their FPBcirc values in HeLa cells. Pearson’s correlation coefficient (r) is shown.

**Extended Data Fig. 5.**
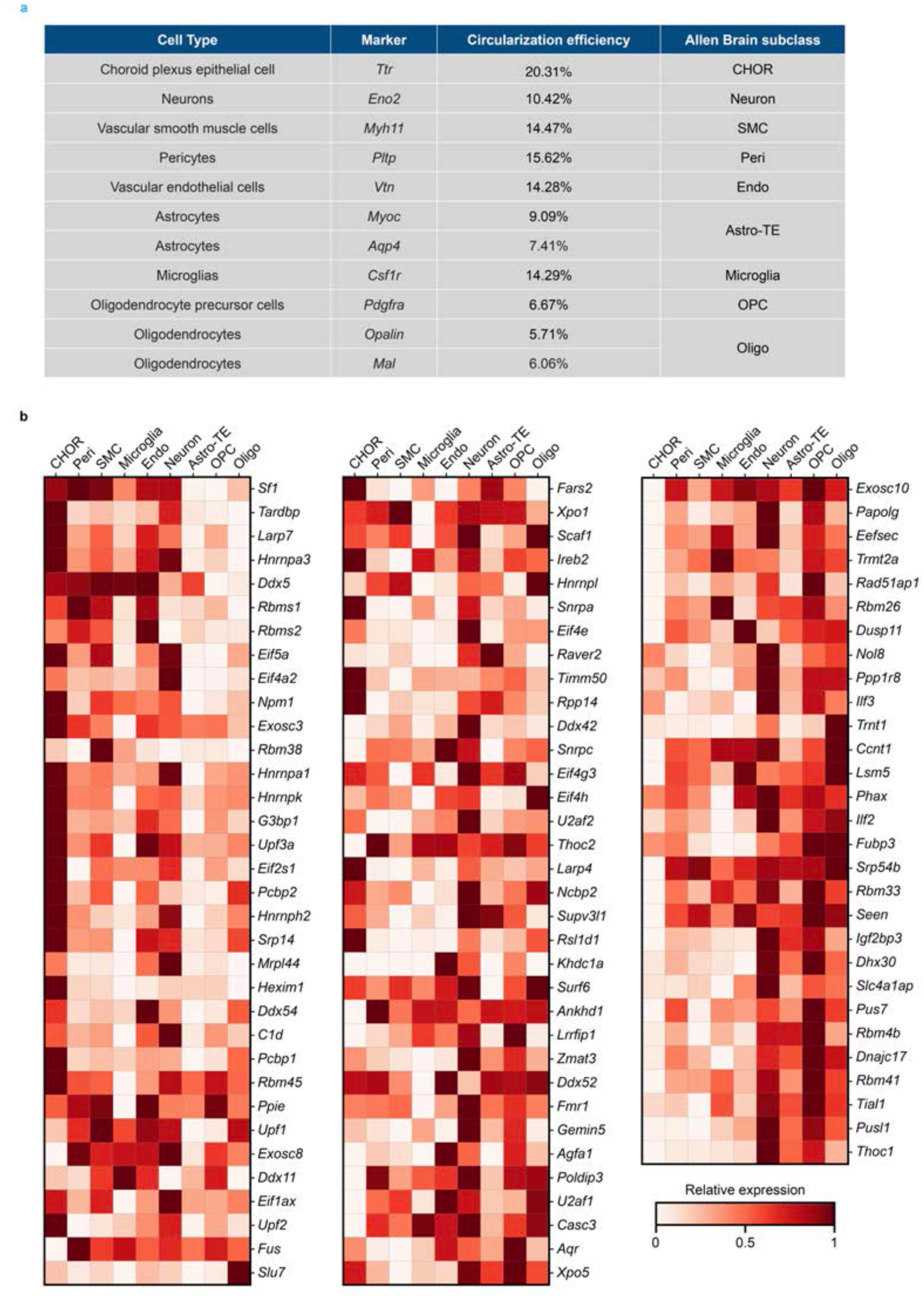
RBP analysis of hippocampus single cell sequencing. **a,** Table of RNA markers and Allen Brain subclasses used to identify major cell types in the mouse hippocampus, with the *circPOLR2A(9,10)* circularization efficiencies. **b,** Heatmap showing expression of circRNA-regulating RBPs across hippocampal cell types from a prior genome-wide screen^46^.

**Extended Data Fig. 6.**
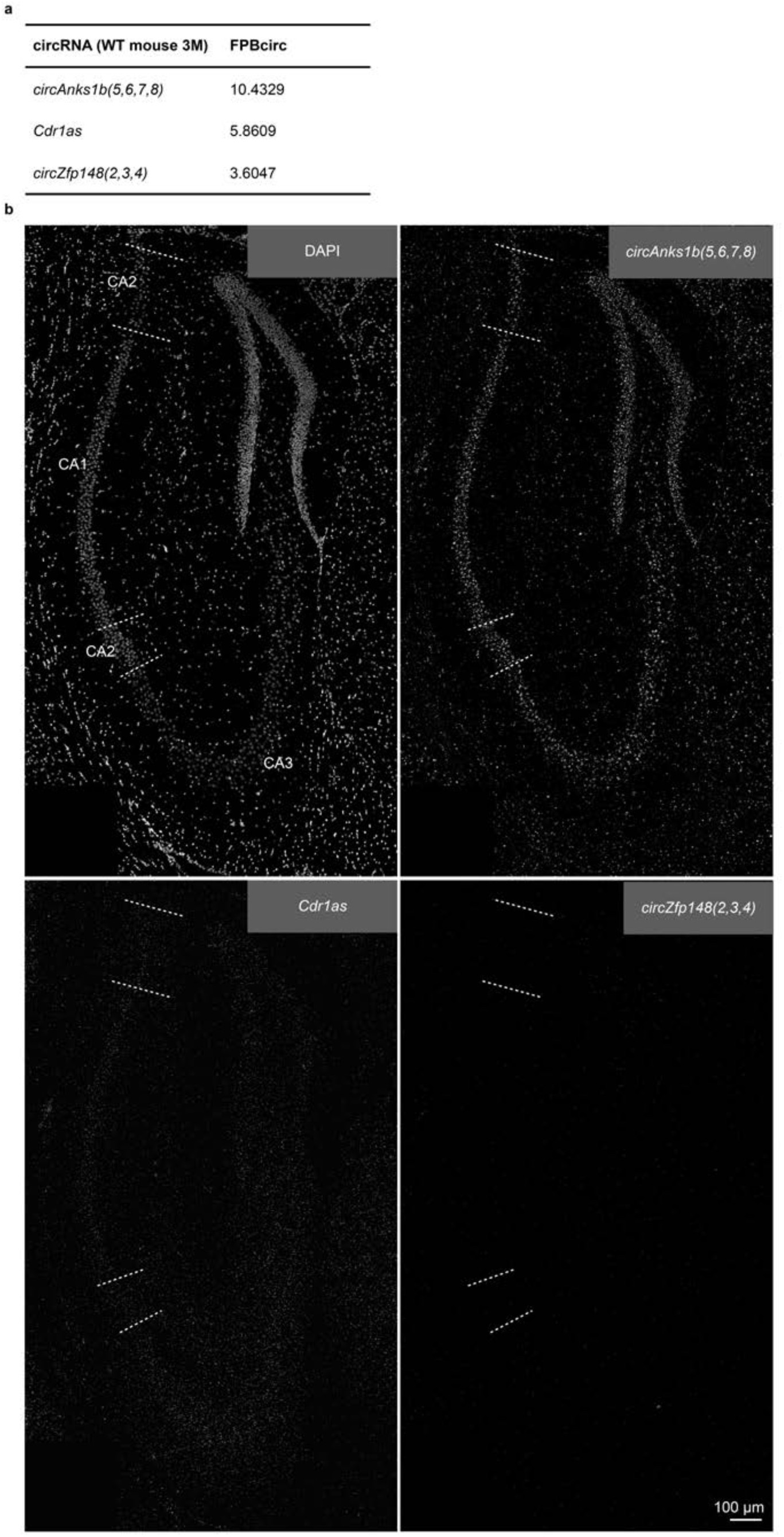
Endogenous circRNAs for *in vivo* imaging. **a,** Table summarizing the FPB_circ_ values of *circZfp148(2,3,4)*, *Cdr1as* and *circAnks1b(5,6,7,8)* in WT mouse (3M) hippocampus. **b,** Single-channel grayscale images of the image shown in Fig. 5a. Subregions CA1, CA2, and CA3 are demarcated by white dashed lines.

**Extended Data Fig. 7.**
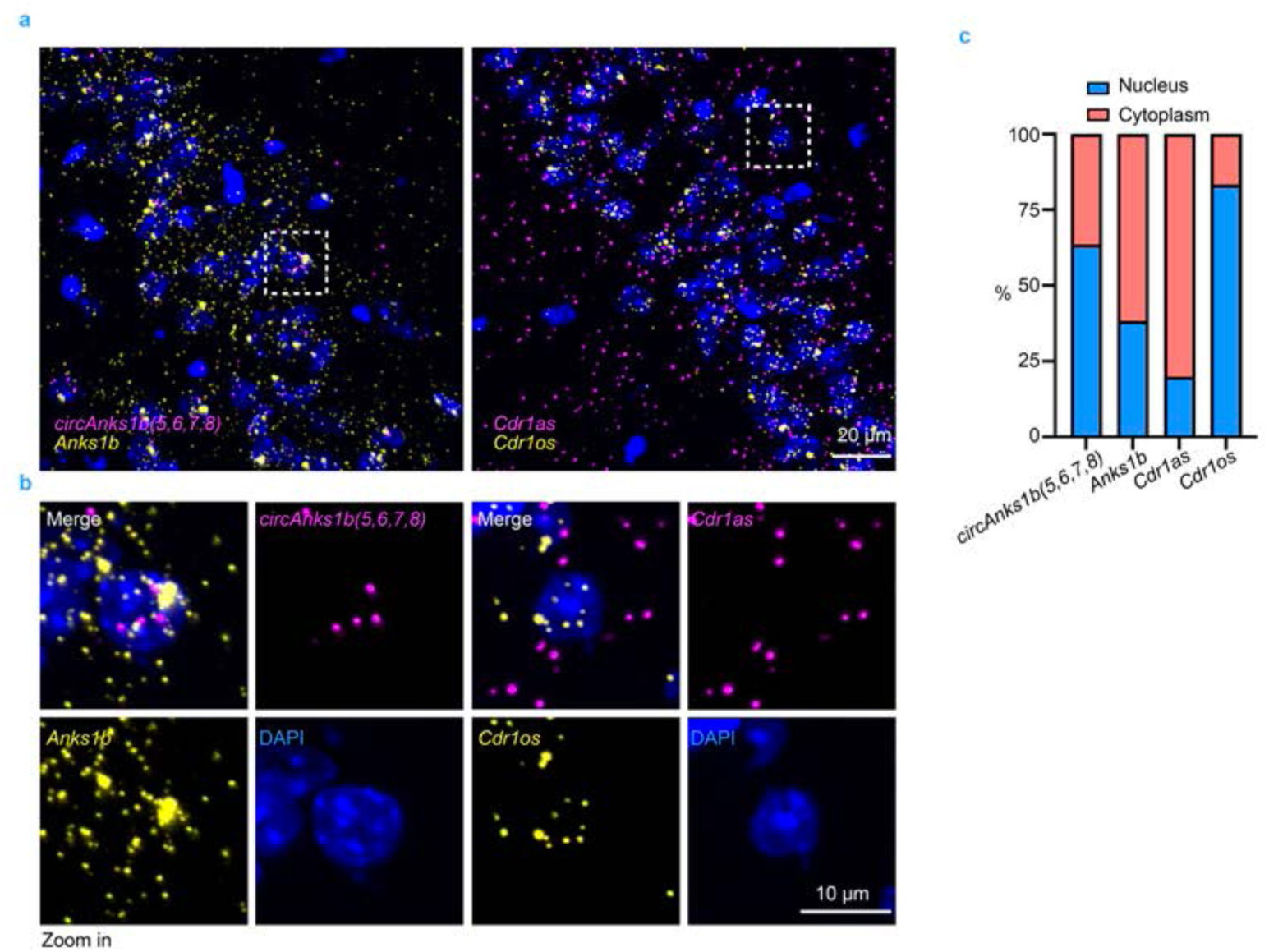
Simultaneous imaging of linear/circular *Anks1b* and *Cdr1as*. **a,** Representative images of simultaneous imaging of linear and circular *Anks1b*, and linear and circular *Cdr1as* in WT mouse (3-month-old) hippocampus. The ROI indicated by white dashed squares is shown at higher magnification in **b**. Magenta, circular RNA signals. Yellow, linear RNA signals. **b,** Higher-magnification view of the ROI from **a**, revealing distinct subcellular localization patterns of linear and circular *Anks1b* and *Cdr1as*. **c,** The percentage of linear and circular transcript subcellular localization in the cytoplasm versus the nucleus. Data were analyzed from 75 cells (*Anks1b*) and 98 cells (*Cdr1as*) shown in **a**. *Anks1b* showed distribution in the cytoplasm while *Cdr1os* exhibited dominant nuclear enrichment.

**Extended Data Fig. 8.**
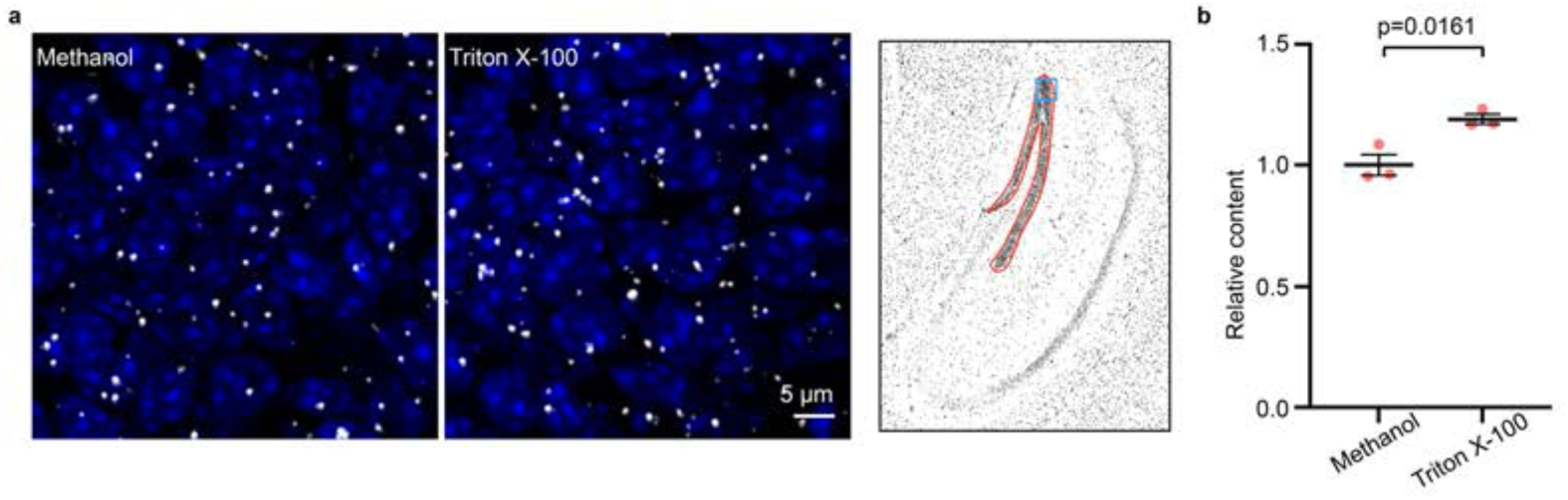
Triton X-100 permeabilization yields a slightly better detection. **a,** Representative abcFISH images of permeabilization optimization for circRNA and protein imaging *in vivo*. The brain sections of KI-mouse were permeabilized with either cold methanol for 1 hour or 0.5% Triton X-100 for 15 min. The images showed partial GrDG shown in blue square on the right panel. White, *circPOLR2A(9,10)* signals. **b,** Statistics of relative *circPOLR2A(9,10)* signal number permeabilized with different reagents shown in **a.** More *circPOLR2A(9,10*) signals were detected in Triton X-100 permeabilized sections. Signal number was calculated within the hippocampal ROI outlined in red in the right panel of **a**. Data represent mean ± s.e.m.; n = 3 brain sections per group; *P* values were calculated using two-tailed Student’s t-test.

